# Combining Scalable Organ Chip Platform with Deep Learning-Based Imaging Analysis for Cancer Therapeutic Screening

**DOI:** 10.1101/2024.06.10.598272

**Authors:** Yu-Chieh Yuan, Beibei Xu, Jenna McCormack, XuHai Huang, Jingzhe Ma, Thomas Marshall, Yacong Sun, Hardeep Singh, Alyssa Fanelli, Paige Gilbride, Xiaohua Qian, Zhiyong Xie, Longlong Si, Xin Xie, Haiqing Bai

## Abstract

Functional precision oncology represents an emerging approach to genomic approaches by testing treatment options directly on patient-derived models. Current assays, including the use of patient-derived xenograft (PDX) and patient-derived organoid (PDOs), faces major barriers in clinical use due to technical challenges, such as standardization, cost, assay time, scalability, and faithful mimicry of patient tumor microenvironment (TME). Here, we introduce an Organ Chip (OC) device constructed entirely from thermoplastic materials, free of porous membrane or other barrier structures, and optimized for high-content imaging (HCI). This automation-compatible device supports tissue-specific extracellular matrices and coculture for a wide spectrum of organ and disease types, including the TME. As a proof-of-concept, we demonstrate the growth of pancreatic, lung, and colon cancer cell lines and primary lung cancer cells and the testing of cancer drugs in these models. HCI-based phenotypic profiling enabled accurate quantification of drug response, with better performance than traditional biochemical assays. Moreover, we developed a deep-learning method for assessing drug responses using bright field images. The integration of a low-cost, scalable, and faithful OC models with automatic high-content image analysis represents a significant stride towards functional precision oncology and cancer drug discovery.

## Introduction

The passage of the FDA Modernization Act 2.0 has catalyzed the integration of advanced in vitro models during drug development^1^, notably in the challenging field of oncology, which is notorious for its high attrition rates^2^. Meanwhile, patient-derived models represent an emerging platform for unraveling the complexity of cancer and tailoring treatment strategies based on lab measured drug response from individual patient’s own tumor cells. Such functional precision oncology assays represent a complementary to genomics-based approaches and may be more widely applicable to different cancer types, drugs or drug combinations ^3^.

Among various PDMs, patient-derived xenografts (PDXs) have been widely adopted due to their ability to maintain the three-dimensional (3D) structure and cellular heterogeneity of human tumors within a living organism. While PDXs offer valuable insights into cancer biology and are commonly used in nonclinical drug efficacy assessment, they are limited by high costs, ethical concerns, and a lack of immune system representation due to their reliance on immunodeficient murine hosts^3^. In contrast, patient-derived organoids (PDOs) represent a more accessible model, allowing for the culture and expansion of tumor cells that retain their original architecture and genetic composition in patients^4^. Nevertheless, the inability of PDO models to mimic the TME with high fidelity, such as the missing of stromal and vascular components, may impede a full understanding of tumor dynamics and compromise the predictive accuracy for determining drug sensitivities with these models^3,5^. Additional bottlenecks impeding their use to guide treatment decision-making include a suboptimal success rate of PDO generation due to insufficient number of viable tumor cells, a lack of standardized assay and predefined threshold to ensure reproducibility and standardization, and a lengthy testing duration from sample collection to report generation (typically exceeding 2 weeks)^3,5^.

By using engineering strategies, such as microfabrication, to create geometries, perfusable channels, and dynamic flows, the Organ Chip (OC) technology enriches the complexity and improves the controllability over other in-vitro models for cell culture, including PDOs. OC holds the promise to simulate the microarchitecture and functions of human tissues and organs with greater human-relevance and translational potential, offering a wide spectrum of applications in studying disease pathophysiology, assessing drug toxicity and efficacy, and advancing personalized medicine^6^. Cancer Chips based on commercial microfluidic devices or laboratory prototypes have been constructed to study pathogenesis across all stages of cancer and adopted for testing anti-cancer therapies^7^. Despite their potential, these systems face challenges in standardization and scalability, and are associated with high costs, which hinder their wider usage beyond research settings.

Here, we introduce a scalable cancer OC platform that marries high-throughput capabilities with advanced imaging-based phenotypic analysis. In a proof-of-concept study, we demonstrate the potential of our platform in drug sensitivity testing for different cancer types. Our platform is primed for clinical drug testing in cancer patients, as well as for preclinical high throughput cancer drug screening.

## Results

### Development of a scalable OC device

A desirable microfluidic device for functional oncology requires a physiologically relevant cell culture system with advanced microfluidics for coculture. This is critical to accurately mimic the complexed TME and to evaluate the effectiveness of various therapeutics.

The system should also adopt biocompatible materials for scalability and cost. To address these needs, we designed a fully thermoplastic microfluidic device, which we termed OC-Plex, that are injection molded and capped with a similar material with thermal bonding, a scalable manufacturing method. Our patented microstructure designs, combined with surface modification techniques, enable a barrierless confinement of hydrogel within two-microchannel microchips (**Fig. 1A**).

**Figure 1.**
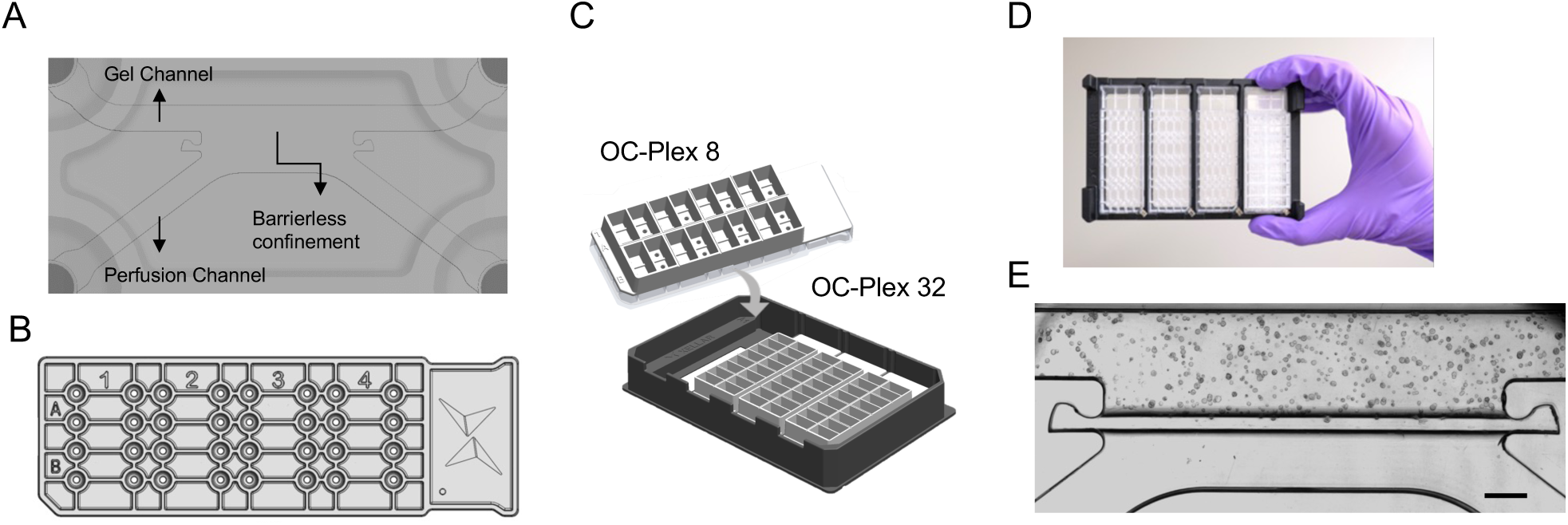
OC-Plex Organ Chip Platforms. **A**) Diagram of the channel designs showing a gel channel and abutting perfusion channel. **B**) CAD drawing of OC-Plex 8. **C**) Each device contains 8 chips; 4 devices can be placed in a stainless-steel carrier for handling or imaging. **D**) Photograph of the OC-Plex 32 product. **E**) Representative imagines showing cells cultured in the device. Scale bar: 500 μm.

Cells can be embedded in the gel (Matrigel, collagen I, or other synthetic hydrogels) and compartmentalized in the gel channel to form a 3D environment with tunable ECM compositions to match in-vivo mechano-properties (**Supplementary Fig. 1**), while the adjacent channel (perfusion channel) facilitates the flow of culture medium. This design permits the establishment of physiologically relevant gradients of nutrients and growth factors across the two channels (**Supplementary Fig. 2**). The OC-Plex supports 8 independent microfluidic chips (**Fig. 1B**) wherein each chip the gel channel and the perfusion channel can be loaded with cell laden hydrogel and cell growth medium, respectively ((**Fig. 1E**). The perfusion channel can be seeded with endothelial cells to vascularize the channel for increased model complexity (**Supplementary Fig. 3**).

The gel and perfusion channels, both with a width of 500 µm and a height of 250 µm converge at the confinement features to create a 3-mm long and 250-µm deep barrier-less culture interface. The microchannels are interfaced with ports spaced in SBS format, enabling automation compatibility. The 8 individual microfluidic chips are multiplexed to form one device, OC-Plex 8, and four OC-Plex 8 devices are further multiplexed with a carrier to form the OC-Plex 32 system (**Fig. 1 C and D**). The OC-Plex is designed to operate in a high-throughput or a low throughput modality for drug screening or device optimization and model research and development, respectively. In the research and development mode, individual devices can be used and coupled with a peristaltic pump for constant flow. While in the high throughput mode, devices can be coupled with an injection molded PS reservoir component to enable gravity driven flow on a rocker. Unlike other organ chip products that employ glass or inert polymers, such as polydimethylsiloxane (PDMS), our device is constructed entirely from thermoplastic materials, making it suitable for mass production with low production costs. Furthermore, our device features low autofluorescence and excellent surface flatness, amenable to imaging-based assays,

### Feasibility of cell growth on chip

To first demonstrate the feasibility of OC-Plex for establishing robust and reproducible cancer OOC models, we developed a pancreatic ductal adenocarcinoma (PDAC) on-chip model using the BxPC-3 cell line. BxPC-3 were embedded in 4 mg/ml Collagen I solution as single cells, loaded into OC-Plex 8 devices in the ECM channel and subsequently cultured for 8 days (**Fig. 2A**). An image analysis pipeline was established to perform image segmentation and feature extraction from Z-projections of the serial Z-stacks of bright field (BF) images (**Fig. 2B**). Feature profiling reveals a consistent increase in the average size of the object (**Fig. 2C top**), as well as the sum area of all objects in a chip during the entire culture duration (**Fig. 2C bottom**). The robustness and reproducibility of the model is demonstrated by consistent growth patterns in three independent experiments (**Fig. 2C**). Additional testing of different seeding densities ranging from 0.3 E+06 cells/mL to 5E+06 cells/mL, corresponding to 300 to 5,000 cells per chip, revealed a diverse phenotypic change over a course of 15 days, including cell aggregate formation, fusion, fission, and migration (**Supplementary Fig. 4A and 4B**).

**Figure 2.**
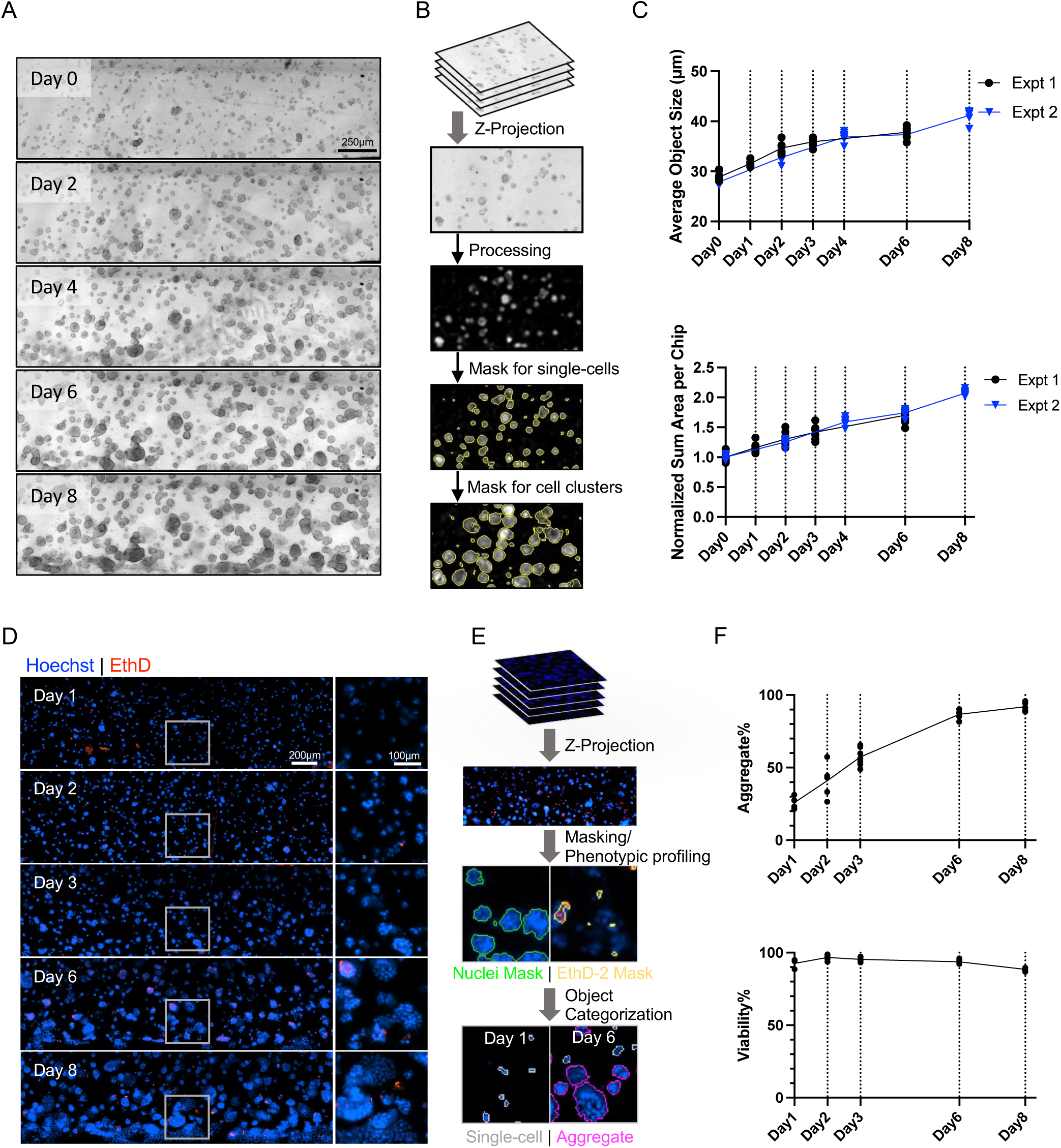
Device supports cancer cell growth. **A)** Representative brightfield images showing single-cell embedded BxPC-3 growing for 8 days in the same OC-Plex chip. **B)** Schematic illustrating analysis strategy for longitudinal imaging to quantify cell growth over time. Brightfield Z-stack images capturing the entire volume of the chips were compressed into Z-projections for image processing and analysis. Each single-cell or cell cluster is segmented as an object, indicated by a yellow contour. **C)** Longitudinal brightfield image analysis results for BxPC-3 growth in OC-Plex. Three experiments were conducted separately, each comprising at least four data points. For the top graph, object size is the average of the object width and length, with each dot representing the mean value per chip. The bottom graph shows the sum area of all objects per chip normalized to the value of day 0 to account for seeding density differences. **D)** Representative chip images of cells stained with nuclear marker (Hoechst33342) and dead cell marker (EthD-2) at various culture time points, demonstrating aggregate formation and minimal cytotoxicity over an 8-day culture period. The squared images in the right column represent the zoomed-in views of the corresponding gray panes in the same row. **E)** Schematic outlining the image analysis strategy for Hoechst33342 and EthD-2-stained z-projection images. Cells are segmented based on Hoechst 33342 staining and further categorized into single-cells (masked in gray) or aggregates (masked in magenta) based on their sizes. Area of dead cells (marked in yellow) is identified by EthD-2 signal. Viable area is quantified by Hoechst33342-stained area minus the dead area. **F)** Image analysis results for Hoechst33342- and EthD-2-stained images with each dot in the graphs representing a chip. The proportion of aggregates to the total cells in each chip increases over time while the cells remain viable.

To evaluate cell viability, Hoechst33342 and Ethidium Homodimer-2 (EthD-2) staining was performed for total cells and cytotoxic cells, respectively (**Fig. 2D**). A fluorescence imaging analysis workflow was developed to segment dead and viable cells and generate imaging masks (**Fig. 2E**). Subsequent analysis at the object level showed that BxPC-3 cells maintained over 95% viability throughout the culture period (**Fig. 2F**), with the percentage of aggregates among total cells on chip increases over time, reaching 100% at day 8 (**Fig. 2F**).

Similarly, aggregate formation was observed for the lung cancer cell line A549 when 100% Basement Membrane Extract (BME) was used to as the embedding hydrogel on chip (**Supplementary Fig. 5A** and **5B**). In addition, cells maintained >90% viability when cultured statically or dynamically with a shaker, despite a difference in the percentage of aggregates under the two conditions (**Supplementary Fig. 5C-E**). Lastly, OC-plex device also supports cell growth and spheroid formation for colon cancer cell lines HCT116 and HT-29 (**Supplementary Fig. 6**). Taken together, these data demonstrated that our device supports the cell growth, spheroid formation, and long-term viability for a wide range of cancer types with different types of extracellular matrix and under different modes of medium perfusion.

### HCI assay development for assessing drug response

Reliable and sensitive readouts are critical for any high-throughput drug screenings. As such, we sought out to develop a workflow for evaluating drug effect in our OC-Plex cancer models using HCI analysis (**Fig. 3A**). BxPC-3 were cultured on chip for 24 hours and exposed to cancer drugs with 0.1% DMSO as negative controls. After 48 hours of drug treatment, viability staining was conducted and whole-chip images were acquired using an automatic microscope.

**Figure 3.**
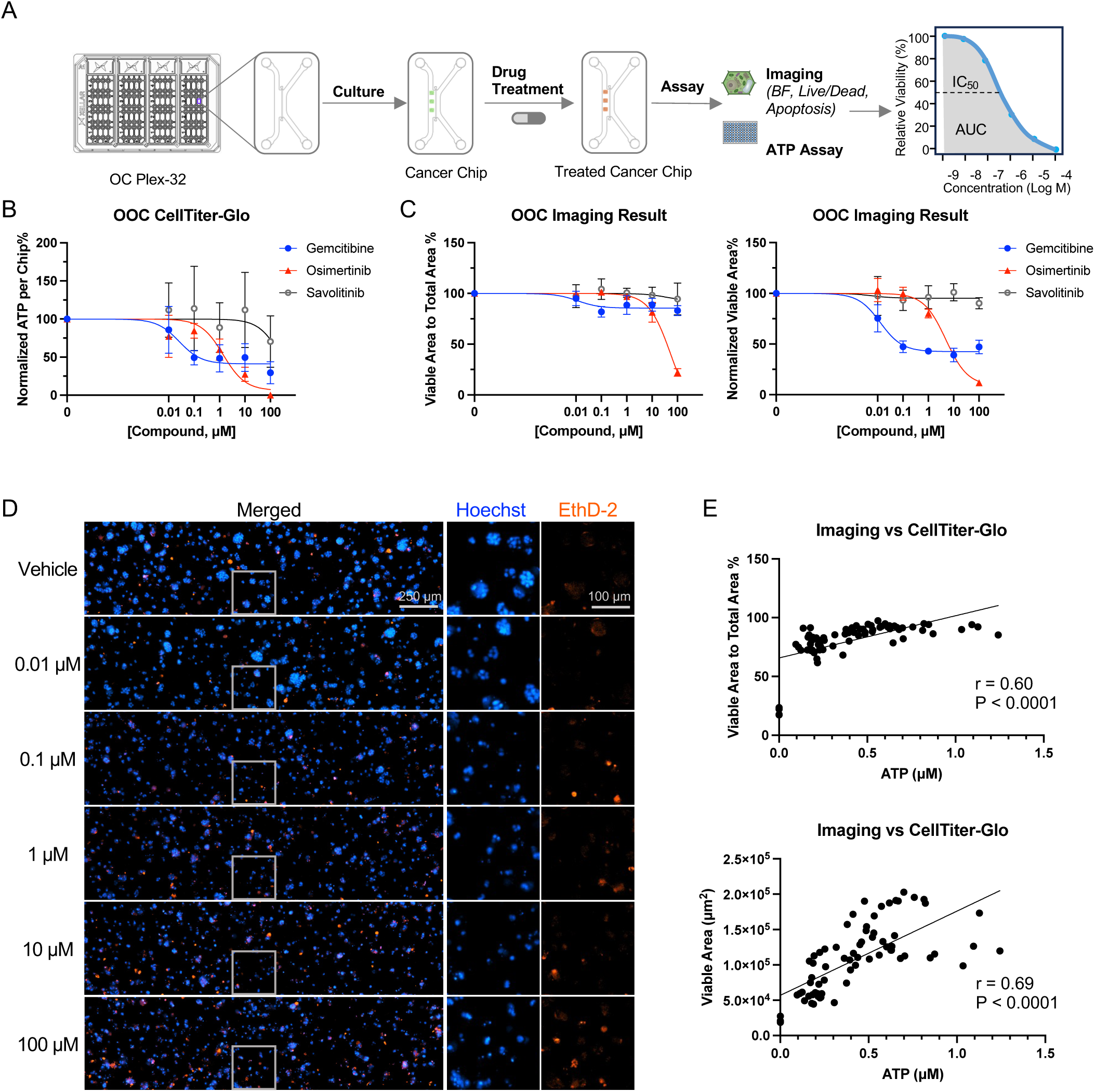
Image analysis enables accurate quantification of drug response. **A)** Schematic of a standardized workflow for high-throughput screening on OC Plex-32 cancer models to measure drug effect **B, C)** Viability results for chips treated with Gemcitabine, Gefitinib, and Osimertinib for 48 hours **B)** by CellTiter-Glo ATP assay and **C)** by imaging assay (n=4). Imaging metrics utilized for dose response curves establishment were selected based on the Z’-factor ranking. The left graph shows the ratio of the area of viable cells to the area of all cells per chip. The right graphs demonstrate the area of viable cells. Data were normalized to the value of the vehicle controls in all graphs in **B** and **C**. **D)** Representative images of chips treated with Osimertinib at 0.01-100 μM for 48 hours and then stained with Hoechst33342 and EthD-2 for viability assessment. The right two image columns are showing the zoomed-in view of the region in the corresponding gray panes in the same row **E)** Correlation graphs comparing imaging results to CellTiter-Glo result for drug effect evaluation. Each dot represents one chip.

To validate the imaging results, we also performed CellTiter-Glo (CTG) assay, the most widely used assay for assessing therapeutic response in 3D cancer models by measuring the amount of cellular ATP^8^. Importantly, we confirmed that the viability staining and CTG assay can be multiplexed, thus allowing the comparison of the two readouts from the same set of devices (**Supplementary Fig. 7A, 7B**).

To establish the HCI assay, we selected three cancer drugs: gemcitabine, gefitinib and osimertinib. Gemcitabine is the standard first-line therapy for pancreatic cancer while epidermal growth factor receptor (EGFR) inhibitors, such as gefitinib and Osimertinib, are two FDA-approved drugs for non-small-cell lung cancer (NSCLC) that have drawn increased attention in treating pancreatic cancer with EGFR activation^9^. CTG results (**Fig. 3B**) indicated that gemcitabine has a smaller IC_50_, whereas EGFR inhibitors, especially osimertinib, resulted in greater inhibition at high concentrations.

For morphological analysis, a total of 60 imaging metrics consisting intensity, shape, and texture features were ranked based on their Z’-factors, a statistical parameter reflective of both the assay signal dynamic range and data variation^10^ (**Supplementary Fig. 7C**). The two metrics with the highest Z’ scores, namely “Normalized Viable Area%” and “Viable Area to Total Area”, were chosen to generate drug dose-response curves. Compared to results from the CTG assay, these imaging metrics revealed a similar trend in ranking drug sensitivity but with much smaller variations (**Fig. 3C**).

Additionally, we observed that the "Viable Area to Total Area” feature exhibited smaller data variations and lower dose-response dynamics compared to “Normalized Viable Area%” (**Fig. 3D**). While “Viable Area to Total Area” can mitigate variations due to seeding density, it also underestimates drug effects that do not involve cell death. This is exemplified by gemcitabine-induced cell growth inhibition (**Fig. 3D**), wherein higher concentrations lead to smaller aggregate sizes without a significant increase in cytotoxicity (EthD-2 signal). In such cases, the normalized viable area is more sensitive in capturing drug effect. Consistently, we observed a higher correlation between ATP levels and viable areas than between ATP levels and “Viable Area to Total Area” (**Fig. 3E**). In addition, we showed that EthD-2 staining can be combined with other cell death markers, such as Caspase-3/7, to enable even greater sensitivity of detection for predicting drug responses (**Supplementary Fig. 8**).

### Cancer Drug Sensitivity Testing

Having established both biochemical and HCI-based readouts to quantify drug response on-chip, we extended the workflow to testing of more cancer types and assessing the impact of culture duration on drug sensitivity. Specifically, we cultured cancer cells for short-term (1 day) and long-term (6-7 days) periods before treating the chips with cancer drugs for 48 hours, with traditional 2D culture in 96-well plate as controls. We initially focused on BxPC-3 pancreatic cancer cells and tested gemcitabine, gefitinib, osimertinib, and savolitinib (a small molecule inhibitor of c-Met used as a negative control). Analysis of drug IC_50_ and the area under the curve (AUC) using imaging metrics revealed a consistent trend in drug sensitivity. However, a decreased drug effect was observed on chip when dosed at day 7 compared to day 1 dosing or 2D control (**Supplementary Fig. 9**). This trend is also visualized in a heatmap comparing the percentage of inhibition across different models (**Fig. 4A**). Consistent with previous analysis (**Supplementary Fig. 7C**), imaging outperformed CTG assay with higher assay sensitivity and lower chip-to-chip variation.

**Figure 4.**
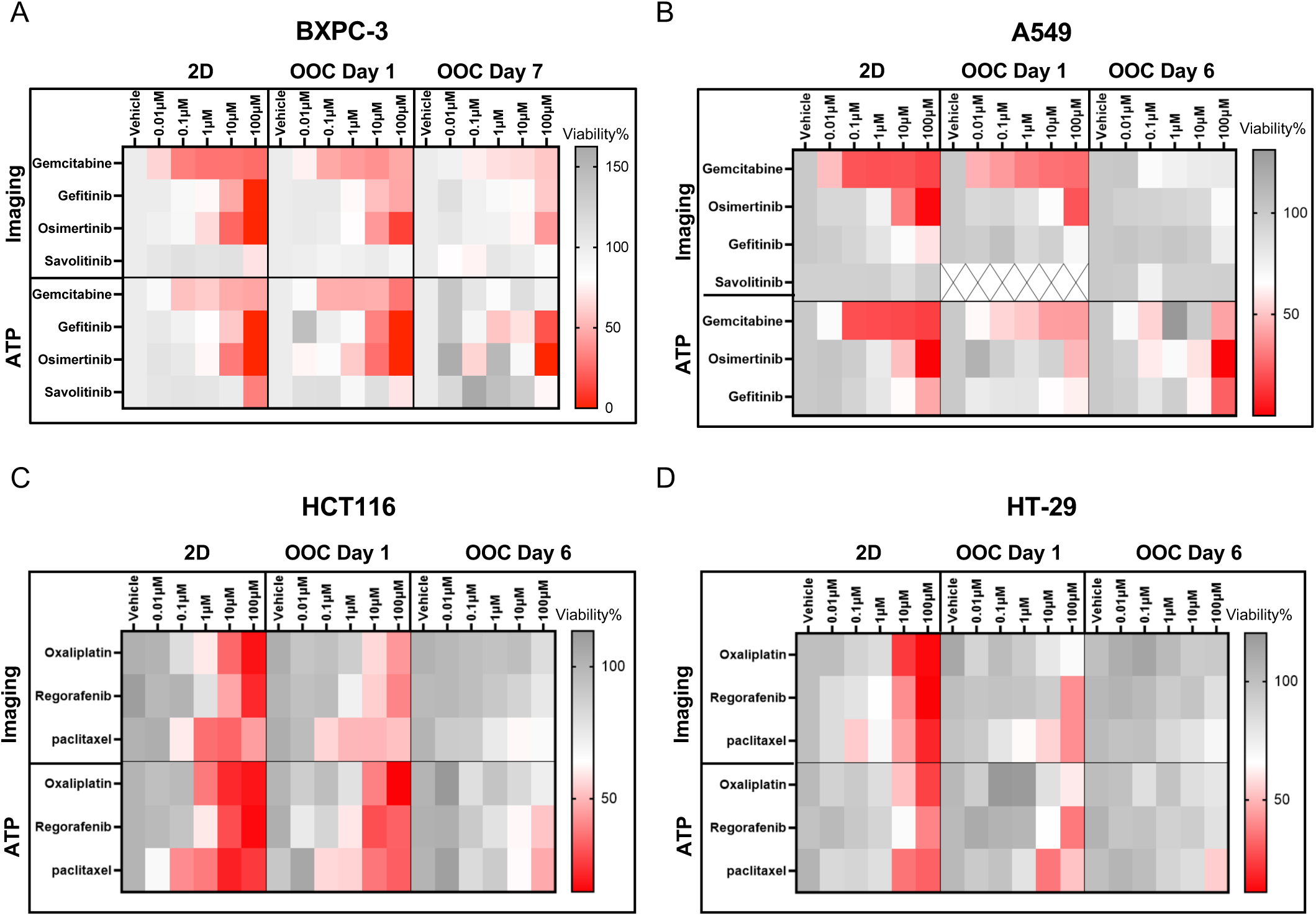
Drug sensitivity testing. **A-D**) Normalized dose-dependent heatmap of cell survival after different drug treatments in 2D and cancer OOC models, survival results were obtained by CellTiter-Glo ATP assay and imaging assay (n=4), respectively. 2D represents the 2D model; OOC Day1 represents the cells were pre-cultured for one day on the OC-Plex device followed by the drug treatment; OOC Day7 represents cells pre-cultured on OC-Plex device for 7 days followed by dosing treatment. Red represents lower cell survival and better drug treatment, and gray represents higher cell survival and worse drug treatment.

We next tested the same set of drugs in the A549 lung cancer model. Imaging analysis confirmed that gemcitabine is the most sensitive drug, followed by gefitinib and osimertinib when drugs were dosed at day 1 after culture on chip (**Supplementary Fig. 10A-B**). Feature analysis confirmed that the same top imaging features selected for BxPC-3 cells can be applied to A549 cells (**Supplementary Fig. 10D-H**). Again, a significantly decreased drug effect was observed on chip when dosed at day 7 compared to day 1 dosing or 2D control (**Fig. 4B**).

For colorectal cancer, we selected representative drugs commonly used in colorectal cancer treatment: oxaliplatin, regorafenib, and paclitaxel^11,12^. These drugs were tested in HCT116 and HT-29 cells in a similar fashion. In general, significant differences in drug inhibition were observed between 2D and on-chip culture (**Fig. 4C-D**): even after one-day culture on chip, both HCT116 and HT-29 cells had significantly reduced sensitivity than 2D control. The sensitivity was further reduced when drugs treatment was initiated six days after culture on chip. These results suggest that the 2D culture model may produce false-positive results while predicting drug response in vitro, and that longer culture duration may offer a closer approximate to in-vivo drug penetration, and thus, tumor responsiveness to drugs. Again, in this model, image analysis results in higher data stability and accuracy compared to the CTG method (**Supplementary Figures 11A,12A**).

### Cancer Drug Sensitivity Testing Using Patient-Derived Cells

To determine whether our cancer chip model could be used for clinical drug testing, we cultured freshly isolated cells from a patient with squamous NSCLC. After 9 days of culture, cell aggregates were observed (**Fig. 5A**). Subsequently, we selected representative therapeutic drugs commonly used in the clinical treatment of lung cancer, including gemcitabine, pemetrexed, paclitaxel, and cisplatin^13^, and treated lung chips with these drugs for 3 days or 6 days prior to culture on-chip for 6 days (**Fig. 5B-C**). In 2D culture, gemcitabine, pemetrexed, and paclitaxel showed comparable efficacy after 72 hours of drug treatment, whereas cells exhibited higher resistance to cisplatin (**Fig. 5D**). However, a different drug sensitivity profile was observed on-chip. Specifically, with a 72-hour dosing regime, cells were most sensitive to paclitaxel, followed by cisplatin, gemcitabine, and pemetrexed. With extended drug dosing of 144 hours, paclitaxel continued to show the strongest efficacy, and the efficacies of gemcitabine and cisplatin increased compared to the 72-hour dosing (**Fig. 5B-C, right**). Pemetrexed efficacy, however, did not significantly increase with prolonged exposure (**Fig. 5B-C**). Unlike the 2D culture results, the lack of efficacy of pemetrexed on-chip aligns with clinical trial findings that patients with squamous NSCLC do not benefit from pemetrexed and face a higher risk of certain adverse effects^14,15^. Therefore, our results demonstrate that our OC-plex device can be used for the culture of patient-derived cancer cells and on-chip drug testing may offer great confidence in predicting drug efficacy in humans compared to 2D models.

**Figure 5.**
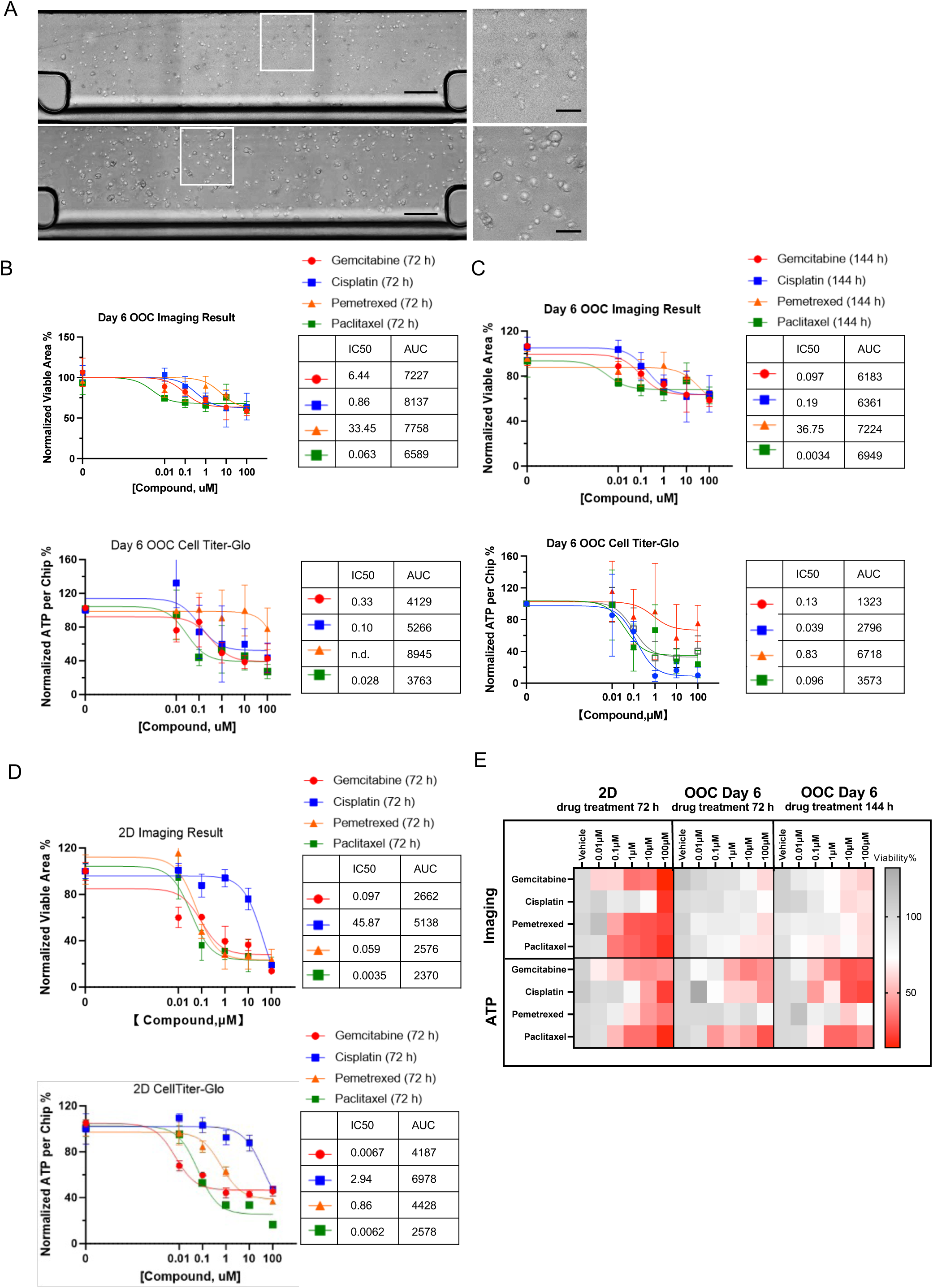
Growth and drug testing of primary cancer cells. **A**) Representative bright field images of primary lung cancer cells cultured on OC-Plex devices for one to nine days. Scale bar: 200 μm. After culturing for six days on OC-Plex devices, primary lung cancer cells were treated with different concentrations of drugs (paclitaxel, gemcitabine, pemetrexed, cisplatin) for 72 h (**B**) or 144 h (**C**), and cell viability was detected by both imaging analysis and CTG assay. **D**) Primary lung cancer cells were treated with different concentrations of drugs (paclitaxel, gemcitabine, pemetrexed, cisplatin) for 72 h under two-dimensional (2D) culture conditions, and cell viability was assessed by imaging analysis and CTG assay. The IC50/AUC (area under the plasma concentration-time curve) was calculated using GraphPad Prism software. **E**) Heatmaps showing the dose-response relationship of cell viability data obtained from panels **A-C**. Red represents lower cell viability, while gray represents higher cell viability.

### A deep learning-based pipeline to enable label-free phenotypic analysis on chip

Fluorescence microscopy enables the visualization of specific (sub)cellular structures through targeted labeling. However, it requires sophisticated equipment and labor-intensive sample preparation. On the other hand, BF imaging captures cellular details without additional preparation and can be performed non-invasively, thus enabling the monitoring of cellular morphology over different time points. Mapping BF to fluorescence images involves training a model to understand the relationship between these imaging modalities concerning the structure of interest, using spatially registered image pairs^16^. To explore whether BF to fluorescence image mapping can be applied to images from our cancer chip, we started by training convolution neural networks with a U-net architecture on paired 2D projected BF and fluorescence images (i.e., DAPI and TRITC) of BxPC3 cells from a training dataset. The model’s performance was then evaluated on a test set, with separate models trained for each fluorescence channel (**Fig. 6A**). The performance was quantified using the Pearson’s correlation coefficient between the extracted features or measurements (e.g., sum intensity, sum area) from the predicted and ground truth images (**Fig. 6B**). The high correlations observed suggest the feasibility of using model-predicted images for feature extraction ((**Fig. 6C**). Features aggregated at the chip level include sum intensity, sum area, and viability.

**Figure 6.**
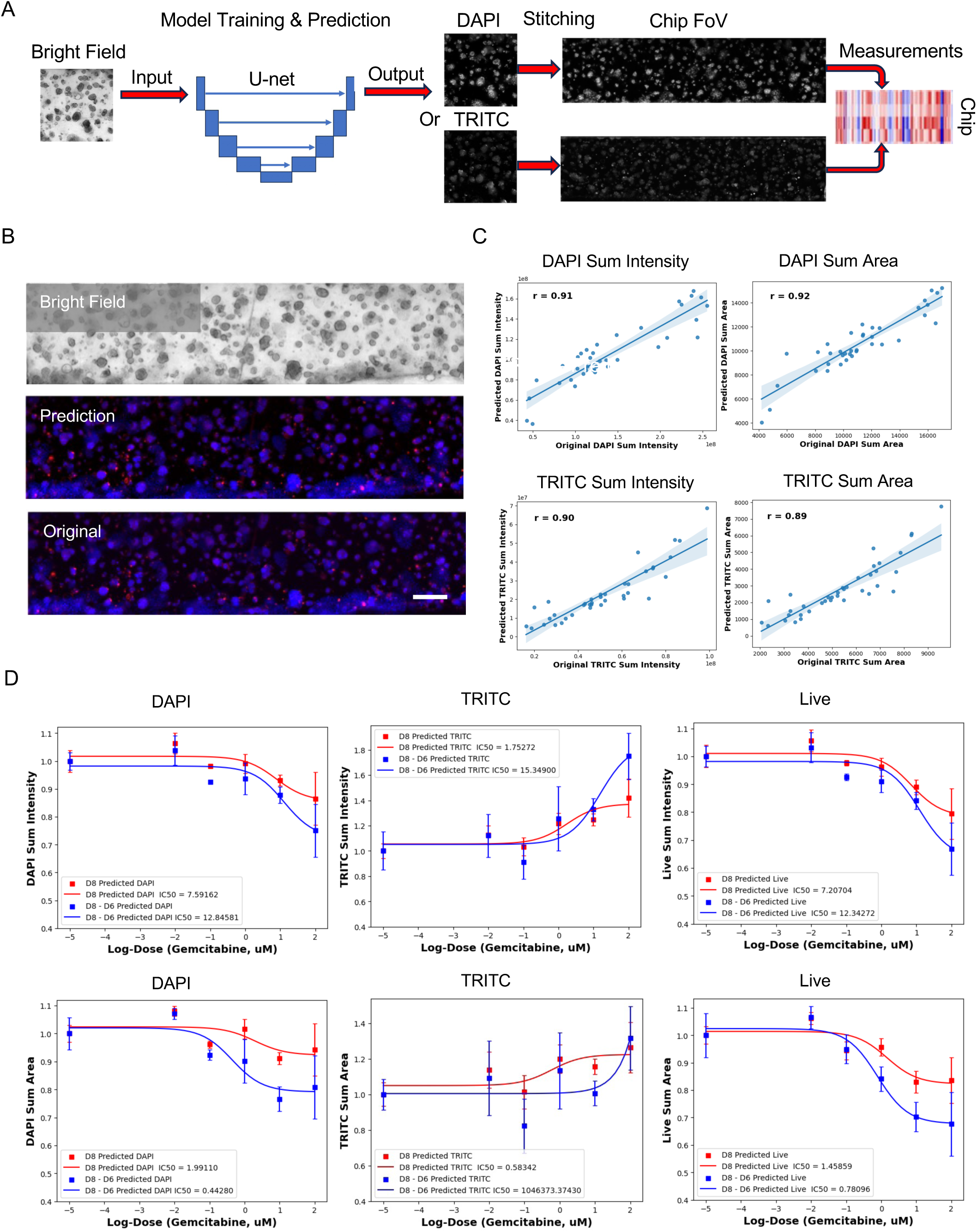
Deep-learning enabled label-free imaging analysis on-chip to quantify drug effect. **A)** Illustration of the analysis workflow. **B**) Examples of original DAPI/TRITC (left) versus model predicted DAPI/TRITC (right) image. **C**) Pearson correlation between experimental measurements and model prediction. (sum intensity/area) extracted from the original DAPI/TRITC images and model predicted DAPI/TRITC on the test dataset. **D**) Dose response curve fitting and IC50 based on sum intensity and sum area measurements. The red and blue squares are the average values over chips at different concentrations of DAPI/TRITC/Live (DAPI - TRITC) at the endpoint (Day 8), and changes from the baseline (i.e., after and before drug treatment), respectively. The red and blue curves are the corresponding best-fit dose response curves.

Next, we examined drug dose responses by fitting dose response curves to chip-level features at different concentrations and calculating IC50 values. Traditionally, drug response evaluation at the endpoint of drug treatment uses fluorescence images taken at the final day (Day 8 in our case). Using in-silico staining, we obtained measurements from the predicted fluorescence images at different timepoints before drug treatment. **Fig. 6D** compares the dose response curves obtained through different methods. The red curve represents data from endpoint images taken on Day 8. Additionally, changes from the baseline are depicted in blue by subtracting baseline measurements taken on Day 6 from the endpoint measurements on Day 8. These comparisons are made using signals from the DAPI channel (left), the TRIC channel (middle), and by masking TRITC-negative regions in the DAPI channel (right). Although all three metrics show dose responses to gemcitabine, changes from the baseline of live cells demonstrate a more sensitive measurement.

In-silico staining also address issues like inhomogeneous staining (**Supplementary Figures 13A**). With predicted fluorescence images at different time points, we could perform longitudinal analysis, which is not feasible with biochemical assay or many staining-based imaging assays. Longitudinal tracking provides insight into cell growth and behavior before and after drug treatment, revealing factors affecting the final drug treatment effect undetectable by endpoint assays (**Supplementary Figures 13B-C**).

## Discussion

In this paper, we introduce a novel organ chip device, OC-Plex, which is modular, scalable, mass-producible, and suitable for high-content imaging. Our device presents several major advantages over existing organ chip platforms. Unlike other platforms, OC-Plex fabrication avoids the use of PDMS, epoxy, and solvent. Instead, it employs injection molding and thermal bonding techniques with polystyrene (PS), similar to commercial cell culture labware, to reduce biocompatibility risks^17^. PS, known for its glass like optical properties^18^, is ideal for high-content imaging and serves as a high moisture barrier while being non-absorptive for small hydrophilic and hydrophobic molecules. While other thermoplastics like COP and COC are used in OOC technology ^19^, PS is preferred for its lower cost and lower glass transitioning temperature, enabling faster turnaround time during thermal bonding. Using injection molding and thermal bonding also ensures low variation in geometric dimensions and high reproducibility compared to conventional PDMS-based OOC, which rely on soft lithography and plasma bonding.

Our proprietary barrier-free confinement design, based on geometric expansion and contraction, not only enables the fully thermoplastic device to be scalable with traditional plastic manufacturing techniques, but also allows unimpeded tissue-tissue interactions between individual channels. This is ideal for applications involving dynamic tissue-tissue interaction, such as the interplay of the tumor cells, vasculature, and the immune cells within the TME ^20^. Additionally, the modular design of the chip layer and reservoir layer offers the flexibility to convert the current single-organ chip to a multi-organ chip configuration in the future by simply modifying the reservoir layer. While our focus in this study was on cancer drug testing, our device has a broad range of potential applications in disease modeling, toxicity testing, and drug screening.

PDOs have not been integrated into prospective clinical decision-making, partially due to a lack of technologies to measure drug response quickly and accurately in clinical settings. By combing high throughput organ chips and HCI analysis, our platform has the potential to address these technical hurdles. To demonstrate the feasibility of our strategy, we performed drug sensitivity screening using both cell lines and patient-derived cancer cells. The reduced drug sensitivity consistently observed across lung, pancreatic, and colon cell lines and patient-derived cells corroborates previous publications and mirrors in vivo findings^21,22^. Possible mechanisms underlying this insensitivity include reduced drug penetration under 3D conditions, altered expression of genes and proteins involved in drug resistance mechanisms (such as increased expression of drug efflux pumps or anti-apoptotic pathways), and the 3D matrix itself acting as a barrier to drug diffsion^23^. However, our results indicating that drug responses from day 1 treatment on-chip approximate those from 2D cultures suggest that the matrix itself is unlikely the reason. Future work is required to decipher the precise molecular mechanisms.

We provide a proof-of-concept for our platform in guiding clinical decision making for treating cancer patients. In our cancer chip model, pemetrexed was not effective against squamous cell carcinoma regardless of the treatment durations (3 or 6 days), despite strong efficacy in traditional 2D testing. This result aligns with clinical recommendations for pemetrexed use in patients with non-squamous non-small cell lung cancer (NSCLC), such as adenocarcinoma and large cell carcinoma, but not for patients with squamous cell carcinoma ^24,25^. Future work involving a larger cohort of patients in a prospective trial is needed to demonstrate the clinical validity of our model.

While ATP-based assays have been widely used in functional precision medicine, our comparative studies suggest that imaging presents a superior readout. Imaging offers several advantages in function precision oncology, including low cost, the ability for multiplexing, providing spatial information, and supporting label free, kinetic analysis^26^. BF imaging excels in tracking dynamic cellular processes over time, facilitating longitudinal studies, whereas fluorescence imaging is limited to providing snapshots of cells at specific instances. Recently, deep learning approaches have been successfully applied to predict fluorescent markers from BF images, known as in-silico labeling/staining ^16,26^. Our approach of using a deep-learning model to achieve BF to fluorescent image translation streamlines the imaging workflow, enhances efficiency, and combines the advantages of both imaging modalities. In-silico labeling/staining can replace the time-consuming sample preparation process with computational techniques and enable longitudinal analysis of predicted fluorescence images. This work pioneers the application of deep learning-based image translation techniques to organ-on-a-chip data, laying the foundation for future endeavors in leveraging state-of-the-art neural network architectures to enhance performance. The deep learning approach holds tremendous potential in saving laborious and costly work for researchers and technicians and offering the opportunity to unveil intricate biological relationships between fluorescent markers.

Taken together, combining our scalable OC-plex Organ Chip Model with imaging and AI analysis addresses major technical barriers to moving functional oncology into clinical practice. While future work is required to increase model complexity by introducing additional components in the TME and to rigorously validate prediction accuracy in clinical trials, our method has the potential to guide individualized cancer care and enable physicians to select specific, effective and safe treatments for their patients.

## Materials and methods

### Chip design and fabrication

The microfluidic channels, OC-Plex 8, and OC-Plex 32 assembly were designed in SOLIDWORKS (Dassault Systems). The injection molding of the OC-Plex was carried out at a third-party manufacturer where a metal injection molding tool was machined with the inverse of the original design. Compared to everyday injection molded plastics, sub-micron surface roughness, micron level deformation, and <1 degree wall draft may change the functionalities of the design geometries; furthermore, achieving micron level flatness is critical in enabling high content imaging. A rigorous design for manufacturing (DFM) process and mold flow (MF) analysis was carried out with the manufacturer to ensure success in injection molding. The molded parts are cleaned with 99.9 2 propanol (Sigma, 34863) and thermal bonded to a piece of die cut 200 µm thick PS films. Due to the strict flatness requirement, proprietary fixturing and bonding strategy was developed to achieve a bond that is robust at elevated temperatures and humidity while maintaining the geometric dimensions specified during injection molding. The bonded part is subsequently functionalized with PEG and sterilized with UV as previously described^27^ to create sterile hydrophilic microchannels. The functional devices are packaged and ready for use.

### Cell Culture

BxPC-3 cells (ATCC, CRL-1687) were cultured in T75 flasks in RPMI-1640 (Gibco, 11835030) supplemented with 10% fetal bovine serum (Gibco, A3160502), 100U/mL of penicillin, and 100μg/mL of streptomycin (Gibco, 15140122) at 37 °C in 5% CO_2_. A549/GFP cells (Cell Biolabs, AKR-209) were cultured in T75 flasks with DMEM, high glucose (Gibco, 11965092). Medium was supplemented with 10% Fetal Bovine Serum (Gibco, A3160502), 0.1 mM MEM Non-Essential Amino Acids (Gibco, 11140050), 1% Penicillin-Streptomycin (Gibco, 15140122) at 37 °C in 5% CO_2_. All cell culture was performed in a humidified incubator at 37 °C, 5% CO_2_. HCT116 (CCTCC, SCSP-5076) and HT-29 cells (CCTCC, SCSP-5032) were cultured in T75 flasks with McCoy’s 5A medium modified (Gibco, 12330031), supplemented with 10% fetal bovine serum (Gibco, A3160502), 100 U/mL of penicillin, and 100 μg/mL of streptomycin (Gibco, 15140122) at 37 °C in 5% CO_2_. All cells used were tested negative for mycoplasma.

The primary lung cancer cells used in this study were isolated from the primary tumor tissues of Lung Squamous Cell Carcinoma (LUSC) patients undergoing surgical treatment. The use of discarded tumor tissues complies with all relevant ethical regulations and has been approved by the Ethics Committee of West China Hospital of Sichuan University (Review No. 1514 in 2023). Briefly, fresh tumor tissues were isolated and promptly transferred to 10 cm dishes. The samples underwent 2-3 washes with PBS and were then finely sliced into 0.1∼1 mm fragments using a scalpel. Subsequently, cells were dissociated utilizing the Tumor Dissociation Kit, human (Miltenyi, 130-095-929). The isolated cells were seeded into T25 flasks containing DMEM, high glucose (Gibco, 11965092) supplemented with 5% fetal bovine serum (Gibco, A3160502), 10 ng/mL bFGF (PeproTech, 100-18B), 10 ng/mL EGF (PeproTech, AF-100-15) and 10 ng/mL IGF (PeproTech, 100-11). The culture medium was refreshed twice weekly until a confluent monolayer was achieved.

### Cell Culture in OC-Plex

The OC-Plex was used for microfluidic cell culture. Rat Tail Collagen Type 1 (R&D Systems, 3447-020-01) prepared at 4 mg/mL with a 1:1:8 ratio of 3.7% (w/v) Sodium Bicarbonate with a pH of 9.5 to 1M HEPES to rat tail collagen, was used as a hydrogel matrix for BxPC3 cell culture and Cultrex Reduced Growth Factor Basement Membrane Extract (BME) (R&D Systems, BME001-05) were used as the hydrogel matrix for A549, HCT116, HT-29 and primary lung cancer cell cultures. Cells were dissociated, pelleted, and resuspended in hydrogel matrix. Hydrogel/cell suspension was kept on ice. Using an electronic single channel pipette, 1.5 μL of hydrogel/cell suspension was seeded into the gel channel. The device was placed in a petri dish with a clean room cloth with 10 mL of molecular grade biology waster and 50 µL of PBS above the ribbed area to prevent gel deformation. Devices were incubated at 37 °C, 5% CO_2_ for 30 minutes (for BME) or 60 minutes (for Collagen-1) to allow polymerization. After incubation, 60 μL of medium was added to the perfusion channel and 20 µL of medium to atop of each gel port to each OC-Plex chip for a total volume of 100 μL per chip. The OC-Plex devices were placed in a humidified incubator and the medium was replaced every 2 – 3 days.

### Drug Treatment

The compounds used in these studies, Gefitinib (Medchem Express, HY-50895), Gemcitabine (MedChem Express, HY-17026), Osimertinib (HY-15722), and Savolitinib (HY-15959), Paclitaxel (HY-B0015), Regorafenib (HY-10331), Pemetrexed (HY-10820) were individually prepared in sterile DMSO as 1000x stocks (0.01, 0.1, 1, 10, 100 mM), aliquoted, and stored at -80 °C. On the day of dosing, each compound was diluted with growth medium at a ratio of 1:1000 to achieve final concentrations of 0.01, 0.1, 1, 10, and 100 µM. Vehicle control was prepared by diluting 100% sterile DMSO to 0.1%. For Cisplatin (HY-17394) and Oxaliplatin (HY-17371), the drugs were prepared as 2 mM master mixes in sterile water and diluted in growth medium to final concentrations of 0.01, 0.1, 1, 10, 100, 200 µM, respectively, before use. Medium was removed completely from each OC-Plex chip or 96-well and replaced with medium containing compound or vehicle for 48 hours prior to endpoint viability assessment.

### Viability Staining

Viability staining solution was prepared by diluting Hoechst 33342 (Thermo Scientific, 62249) by 1:2000 to achieve a final concentration of 10 μM, Ethidium Homodimer-2 (Invitrogen, E3599) by 1:1000 to achieve 1 μM, and optionally, CellEvent^TM^ Caspase-3/7 green detection reagent (Invitrogen, C10423) by 1:400 to achieve 5 μM in complete growth medium. Prior to staining, the medium was removed completely from OC-Plex and 96-well plate and replaced with staining solution. Stain was incubated for 2 hours for OC-Plex and 40 minutes for a 96-well plate in a humidified incubator (37°C, 5% CO_2_) before imaging. For OC-Plex, staining solution was mixed by pipetting up and down in the perfusion channel and ECM ports after 1 hour to enhance staining efficiency.

### Image Acquisition and Processing

Imaging was conducted with an Agilent BioTek Lionheart LX Automated Fluorescent Microscope through a 4x air objective (Olympus Plan Fluorite 0.13NA #1220519). Automated imaging and processing protocols were established using BioTek Gen5 Software (Agilent). Each chip was acquired with two 10%-overlapping fields of view and 19 z-slices (14-μm step size) to encompass the entire ECM channel height. Longitudinal imaging for cell growth motoring was performed in the brightfield channel. Pre-analytical processing steps were applied to the brightfield images, including stitching, 2D projection, digital phase contrast, kinetic frame registration, background subtraction, and smoothing. Live/Dead Imaging was acquired in DAPI channel (Excitation 377/50 nm, Emission 447/60 nm) for Hoechst 33342, TRITC channel (Excitation 556/20 nm, Emission 600/37 nm) for EthD-2, and GFP channel (Excitation 469/35 nm, Emission 525/39 nm) for Caspase-3/7. Fluorescence images for each channel underwent stitching, background subtraction and 2D projection before image analysis.

### Image Analysis

Image analysis was performed using BioTek Gen5 Software (Agilent). Processed brightfield images were utilized for cell/aggregate segmentation to monitor cell growth over time. Each individual cell or cluster of cells was segmented as an object based on its intensity. The changes in mean object area over time of each OC-Plex chip was quantified to represent the average growth of individual cells or clusters. Additionally, the changes in the sum area of all objects in each chip over time were calculated to indicate overall the cell growth the chip-level.

For viability imaging assay, the image analysis strategy involved confining region of interest (ROI) and feature extraction using a fluorescence intensity-based masking process. Individual cells or clusters of cells was segmented as objects by applying a mask using thresholding method based on Hoechst 33342 intensity. Within the object mask, dead cells and apoptotic cells were identified based on EthD-2 intensity and Caspase-3/7 intensity, respectively. Each object was further assessed for a variety of intensity and morphology related metrics. Following the calculation of object morphology and intensity, objects were categorized into “Single-Cell’ and “Aggregate” groups based on their sizes. The viable area was quantified by deducting the EthD-2 mask area from Hoechst33342 mask, and the viability% was quantified by using viable area normalized to the sum area of all cells.

### Z’-Factor Analysis

Z’-Factor was calculated based on the equation provided below:

Z’ = 1 – (3αc+ + αc-)/ |μc+ – μc-|, where αc+ and αc-represent the standard deviations of the positive control and negative control respectively and μc+ and μc-represent the means of the positive control and negative control respectively10. For drug sensitivity testing, the vehicle control (0.1% DMSO) serves as the negative control; 100 μM Osimertinib and 100 μM Gemcitabine serve as the positive controls for BxPC-3 and A549 models, respectively. Z’-factor was calculated using all imaging metrics and CTG.

### Fluorescent tracer penetration assay

Collagen 1 gel was used to access the diffusion rate from the perfusion channel to gel channel. Rhodamine B was used as the tracer molecule for its hydrophilicity and auto fluorescence. A 200 µM of rhodamine B (Sigma, 83689-1G) solution was prepared in 1x PBS and loaded into the perfusion channel. The OC-Plex chip with the gel and rhodamine B was allowed to incubate at room temperature and imaged with brightfield at time 0 min, 2.5 min, 5 min, and 10 min. To increase fidelity, kinetic imaging was carried out to assess diffusion rate with 1 µM and 10 µM of rhodamine B and Cy5-Dextran. A 70 kDa Cy5-Dextran (Sigma, 90718-1G) was added to the experiment to account for a large hydrophilic molecule. The OC-Plex chips was incubated at room temperature for 1 hour and 2 hours for 1 µM solutions and 10 µM solutions, respectively. During the incubation period, fluorescent images were taken every 2 minutes with an Agilent BioTek Lionheart LX Automated Fluorescent Microscope.

### Cell Viability Assay

To quantify cell viability in the OC-Plex device, culture medium was removed from both channels. Subsequently, 100µL of CellTiter-Glo 3D Reagent (Promega, G9681) was aspirated and dispensed into the open media channel and ECM ports. . This process was repeated for all chips in the device. Following this, the device was placed on an orbital shaker at room temperature for 1 hour. Concurrently, a rATP standard (Promega, P1132) was prepared by diluting the 10mM stock 1:1000 in culture medium. 50µL of this medium was added to wells A2-A12 of a white opaque 96-well plate, and 100µL of the diluted rATP standard was placed into well A1. This standard was serially diluted by transferring 50µL from well A1 to well A2 and so on, mixing each well 5-7 times, with well A12 reserved as the blank. After shaking the plate for 5 minutes, it was incubated for 25 minutes. Upon completion of the 1-hour incubation of the devices, the CellTiter-Glo 3D Reagent was aspirated from the devices and transferred to the 96-well plate, with each chip’s reagent in separate wells. Luminescence was read immediately with an integration time of 0.25-1 second per well.

### Model Training, Validation, and Testing

We employed the U-net convolutional neural network due to its demonstrated performance in image-related tasks and relative simplicity (relatively fewer parameters thus easy to train). The deep learning model was trained and validated on a BxPC3 cell line dataset of a total of 756 image pairs for each channel. We split the dataset into training, validation, and test sets with a ratio of 7:2:1. The validation set was used for hyperparameter tuning and the test set was used to evaluate the final model performance.

The Z-projection image has a region of interest of 1050 px X 260 px and was cropped into 224 px X 224 px patches with overlapping regions as the input image to U-net. We used a batch size of 8 and the Adam optimizer with a learning rate of 10-4. The model was trained using an MSE loss over 150 epochs. The model performance was evaluated via the Pearson’s correlation coefficient on extracted features or measurements between predicted fluorescence images and ground truth images.

### Image Feature Extraction and Analysis

We extracted features from the predicted chip-level images after stitching image crops.

The post-processing steps include Gaussian smoothing, artifact removal, thresholding (Two-class Otsu thresholding used for DAPI and Three-class Otsu used for TRITC), and threshold-based segmentation. The sum intensity and sum area are the two most representative measurements of the chip-level properties. We also measured the cell viability using different metrics. The live area/intensity in Figure 2 is defined as the DAPI area/intensity minus the TRITC positive area/intensity within the DAPI mask. The area/intensity-based viability in Figure 3 is the ratio of live area/intensity to DAPI area/intensity.

The longitudinal predictions utilized the relationship between BF and fluorescence images learned at the endpoint images (Day 8) and applied to BF images taken at Days 0, 2, 4, 6. The image features at earlier timepoints were extracted in the same manner as the endpoint. Day 8 – Day 6 properties represent the changes from the baseline due to drug treatment, which alleviate the impact of chip-to-chip variation. The earlier timepoints track the cell behavior before drug treatment.

### Statistical analysis

All experiments were repeated at least two times. Data are displayed as mean values ± standard deviations (SD) unless otherwise noted. Graphing and statistical comparison of the data were performed using GraphPad Prism 9.0. Two-group comparisons were assessed using the two-tailed Student’s t test; comparison of three or more groups were analyzed by one-way ANOVA with Tukey’s multiple comparisons test. *p* values < 0.05 were considered to be statistically significant; * p< 0.05; **p< 0.01; ***< 0.001; n.s., not significant.

### Author Contributions

H.B. and X.X. conceptualized the study and guided experimental designs. H.B. drafted the manuscript with input from all authors. YC.Y., B.X., and J.MC. performed experiments involving the pancreatic cancer, colon cancer, and A549 cells respectively. B.X. performed experiments involving patient-derived lung cancer. XH. H. conducted engineering test of the device. J.M. and Z.X. performed deep-learning based image analysis. All authors contributed to the revision of the manuscript. The final version of manuscript has been approved by all authors.

## Acknowledgment

We would like to thank Dr. Adam J. Bindas who was involved in the prototyping stage of the OC-Plex device.

