## Supplementary Information for "Combining Scalable Organ Chip Platform with Deep Learning-Based Imaging Analysis for Cancer Therapeutic Screening"

**Supplementary Figure 1. Filling success rate for different ECM in OC-Plex device.** The OC-Plex chip is designed to enable filling and confining hydrogel in the gel channel. The filling and compartmentalization of the gel is achieved by the surface modification and geometric design within the microfluidic chip. To demonstrate hydrogel compatibility, 3 concentrations of BME and Collagen-1 were tested. 32 chips were filled with each of the gel in figure 2, where all gel showed >90% success rate.

**Supplementary Figure 2. Permeability assay on Chip. A**) Representative images showing the diffusion of fluorescent dyes from the perfusion channel into the gel channel. **B**) Quantification of the diffusion **C**) First derivative of ECM to medium channel intensity ratio.

**Supplementary Figure 3. Support of coculture on-chip. A)** Devices placed in a stainless steel carrier to allow attachment of cells on the gel interface. **B)** Image analysis results showing consistent cell counts, each dot represents one chip. **C)** Endothelial cell attachment at gel interface after 1-hour incubation in carrier. **D)** Dynamic culture with reservoir and shaker to allow flow. **E)** Chips connected via tubing to peristaltic pump for high shear applications. **F)** Representative image of tubule formation of endothelial cells in the perfusion channel labeled with CellTracker.

**Supplementary Figure 4. Growth of pancreatic cell line on-chip A)** Longitudinal brightfield image analysis results of BxPC-3 growth on OC-Plex32 chips at four seeding densities ranging from $0.31\times10^6$-$5\times10^6$ cells/mL (n=4). The normalization for the left graph is done by dividing the values to the values of day 0 for each chip. **B)** Viability results obtain from CellTiter-Glo assay for various seeding densities. For $5\times10^6$ and $0.31\times10^6$, the viability was assessed after 18-day culture period. For $2.5\times10^6$ and $1.25\times10^6$, viability was assessed after 17-day culture period.

The dead control was induced by adding 0.2% Triton X-100 2-hour prior to adding CellTiter-Glo lysing buffer **C)** Representative brightfield images of single-cell-embedded BxPC-3 growth at different seeding densities over 15-18 days culture period.

**Supplementary Figure 5. Growth of lung cell line on-chip A)** Representative brightfield images showing single-cell embedded A549 growing for 8 days in the same OC-Plex chip. **B)** Longitudinal brightfield image analysis results for A549 growth in OC-Plex, the top graph shows the mean area per object and the bottom graph shows the sum of all objects per chip normalized to the value of day 0 to account for seeding density differences. **C)** Representative chip images of cells staining with nuclear marker (Hoechst33342) and dead cell marker (EthD-2) at day 7 of culture in static with reservoir and dynamic culture showing comparable viability between culture methods. **D)** Image analysis results of viable area for Hoechst33342 and EthD-2 stained images, each dot represents one chip. **E)** The proportion of aggregates to the total cells in each chip at day 7 of culture for different culture methods, the proportion of aggregates increases when cultured with the reservoir attachment.

**Supplementary Figure 6. Growth of colon cell line on-chip.** Representative images of HCT116 and HT-29 cells cultured on the OC-Plex device for one day to eight days, respectively.BF indicates bright field image, and Hoechst/EthD-2 indicates a representative image of viability assessment with Hoechst33342 and EthD-2 staining. Scale bar: 200 μm.

**Supplementary Figure 7. Effect of live/dead staining on ATP assay A)** Comparison of the CellTiter-Glo ATP results with and without Hoechst 33342 and EthD-2 staining addition **B)** ATP results obtained from chips stained with Hoechst 33342 and EthD-2. The chips were seeded at densities ranging from $0.31\times10^6$-$5\times10^6$ cells/mL and imaged for viability prior to the ATP assay.

**Supplementary Figure 8. Combining multiple staining on chip. A)** Representative images showing viability assessment by Hoechst33342, EthD-2, and caspase 3/7 staining after drug treatment. **B**) Schematic of the workflow for segmentation and downstream analysis of the triple-stained fluorescent images. **C**) Normalized Drug dose response curves of BXPC-3 cells by using staining from EthD-2, caspase 3/7, or both. **D**) similar to **C**) but without normalization to total Hoechst area. Scale bar: 200 µm.

**Supplementary Figure 9. Drug Testing in pancreatic cancer chip.** A) Representative images showing viability assessment by Hoechst33342 and EthD-2 staining after treatment with Gemcitabine (**A**), Gefitinib (**B**), Osimertinib (**C**), and Savolitinib (**D**) at indicated concentrations from 0.01 to 100 µM for 48 hours under 2D condition or after on chip culture for 1 day or 7 days. Drug effect on BxPC-3 cell viability were analyzed using CellTiter-Glo ATP assay or imaging detection (n=4). The dose response curves were plotted (**E**). IC50 and AUC (area under the plasma concentration-time curve) were calculated using GraphPad Prism software. All data are presented as the mean ± SD of three replicates.

**Supplementary Figure 10. Drug Testing in lung cancer chip. A-D)** Dose-response curves of A549 cells treated with gemcitabine, osimertinib, or gefitinib after 1 day (**A&B**) or 6 days of culture (**C&D**) and analyzed using CellTitre-Glo (**A&C**) or imaging assay (**B&D**). **E** shows representative images from gemcitabine-treated chips. **F**) feature ranking by Z' score. **G-H**) Correlation between the top 2 features and ATP results.

**Supplementary Figure 11. Drug Testing in colon cancer chip for HCT116.** A) Representative images showing viability assessment by Hoechst33342 and EthD-2 staining after treatment with oxaliplatin (**A**), regorafenib (**B**), and paclitaxel (**C**) at indicated concentrations from 0.01 to 100 μM for 48 hours under 2D condition or after on chip culture for 1 day or 6 days. Drug effect on HCT116 cell viability were analyzed using CellTiter-Glo ATP assay or imaging detection (n=4). The dose response curves were plotted (**D, E**). IC50 and AUC (area under the plasma concentration-time curve) were calculated using GraphPad Prism software. All data are presented as the mean ± SD of three replicates.

**Supplementary Figure 12. Drug Testing in colon cancer chip for HT-29.** A) Representative images showing viability assessment by Hoechst33342 and EthD-2 staining after treatment with oxaliplatin (**A**), regorafenib (**B**), and paclitaxel (**C**) at indicated concentrations from 0.01 to 100 μM for 48 hours under 2D condition or after on chip culture for 1 day or 6 days. Drug effect on HT-29 cell viability were analyzed using CellTiter-Glo ATP assay or imaging detection (n=4). The dose response curves were plotted (**D, E**). IC50 and AUC (area under the plasma concentration-time curve) were calculated using GraphPad Prism software. All data are presented as the mean ± SD of three replicates.

**Supplementary Figure 13. Advantages of combing bright-field imaging and deep learning algorithms for assessing drug responses on chip. A**) Examples of in-silico staining overcome staining homogeneity issues with experimental staining. **B, C**) Longitudinal tracking to measure sum intensity and sum area across DAPI/TRITC/Live channels.

**Supplementary Figure 1**

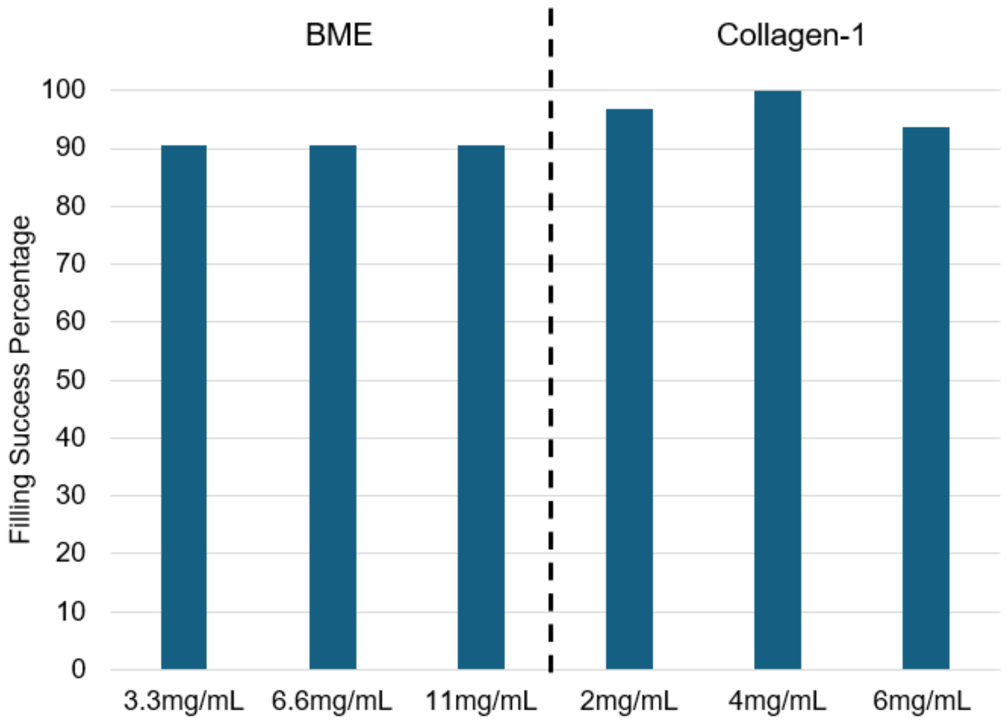

#### Supplementary Figure 2

A

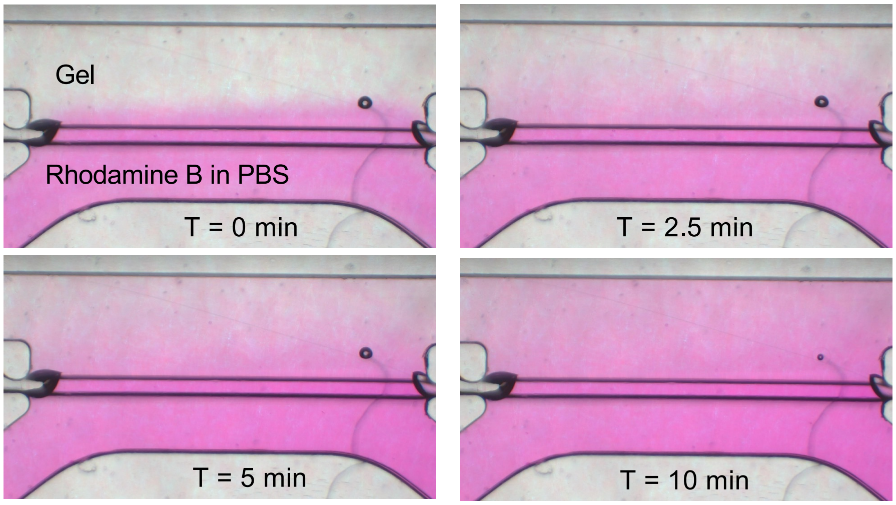


B

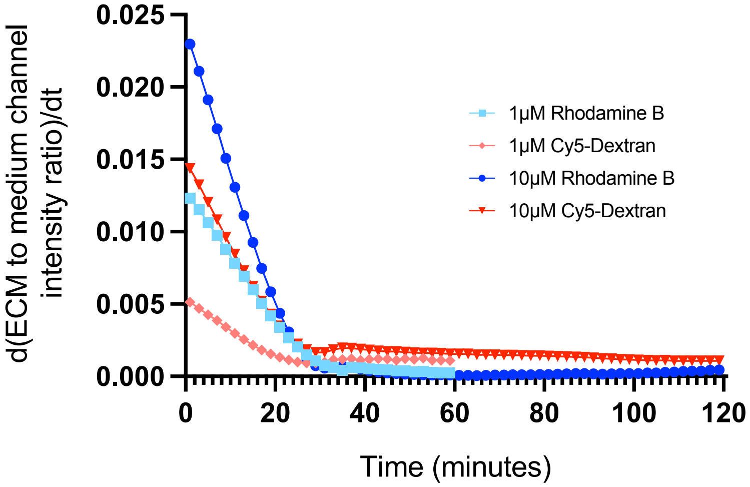

### Supplementary Figure 3

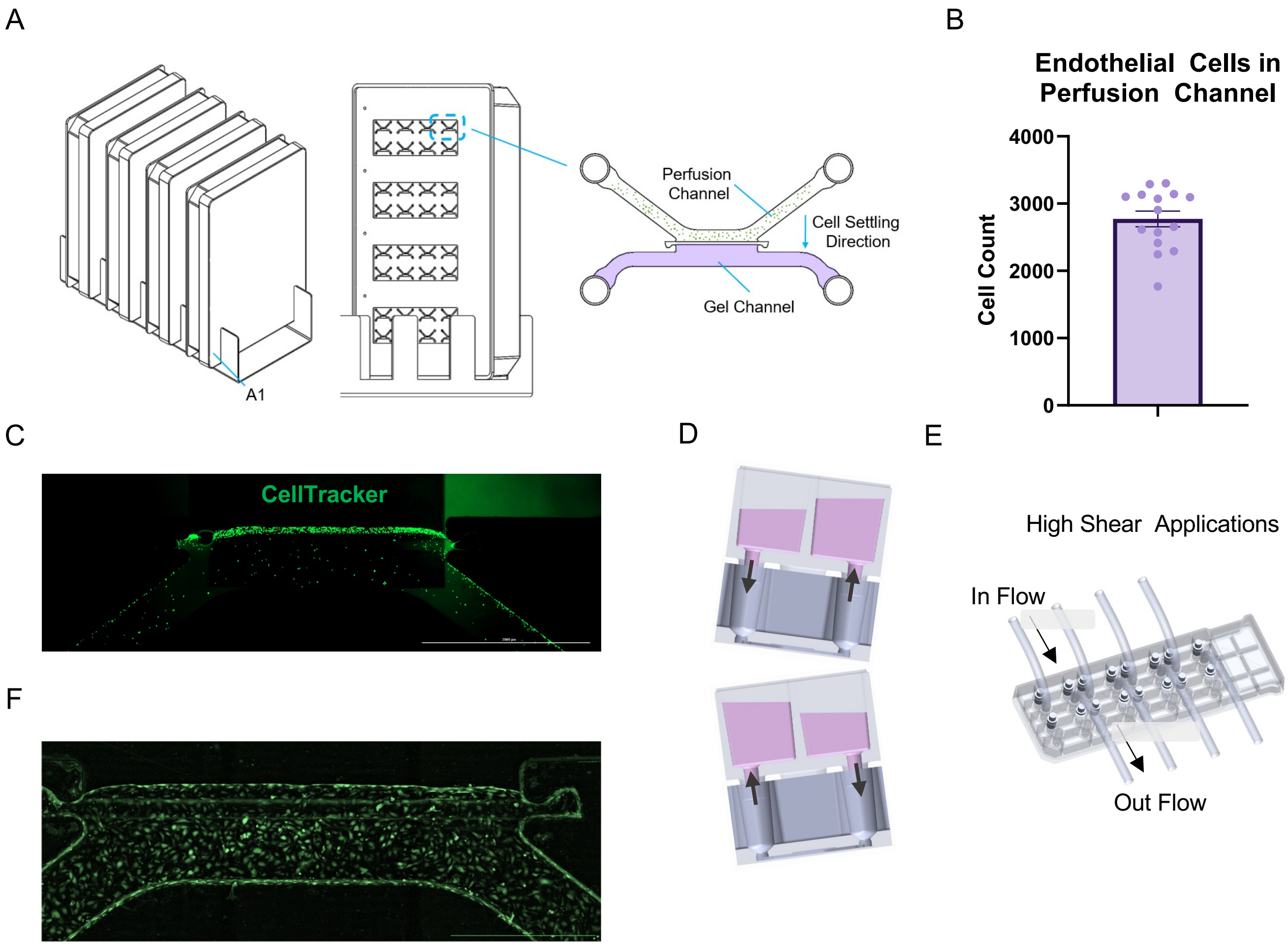

### Supplementary Figure 4

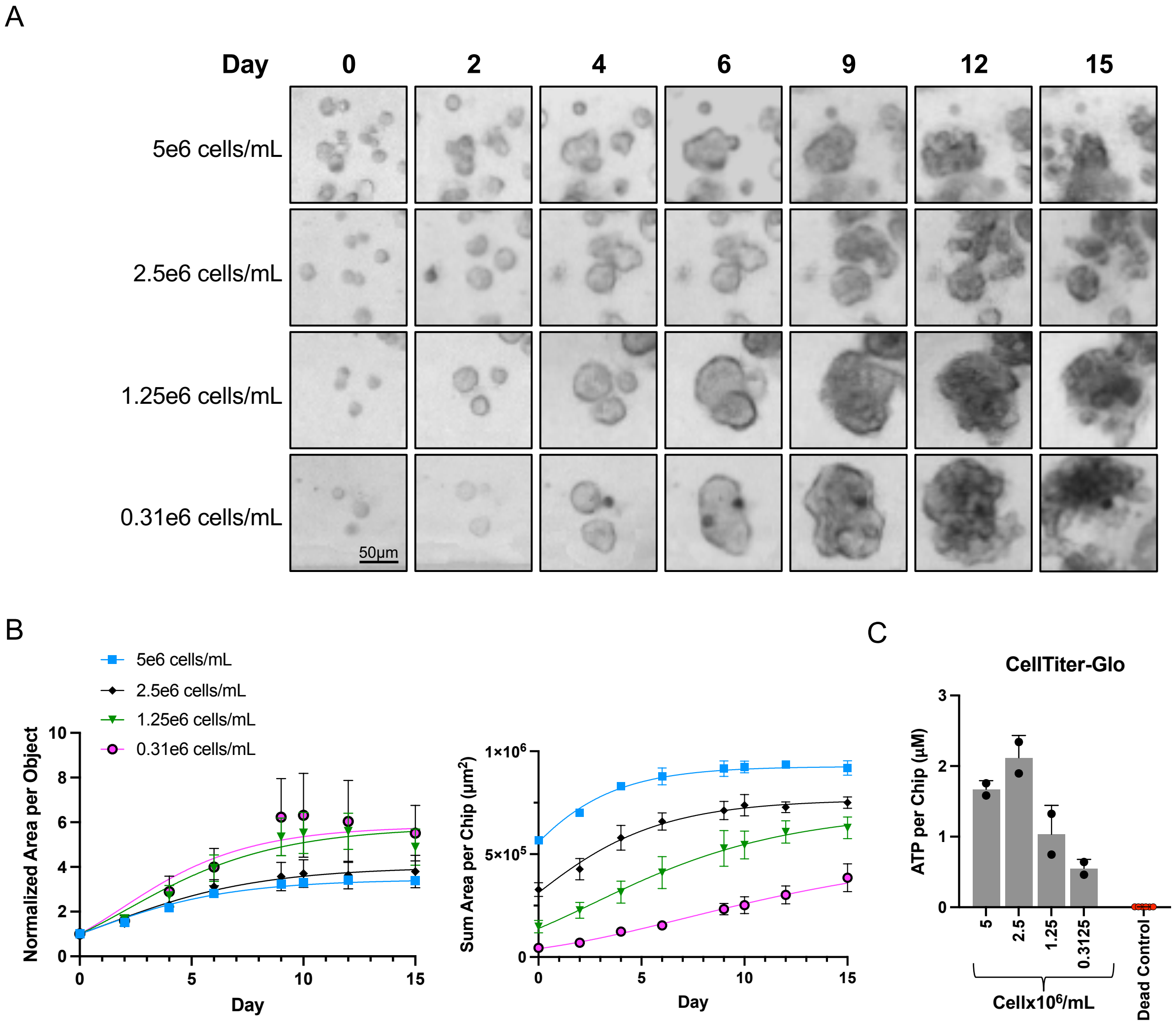

#### Supplementary Figure 5

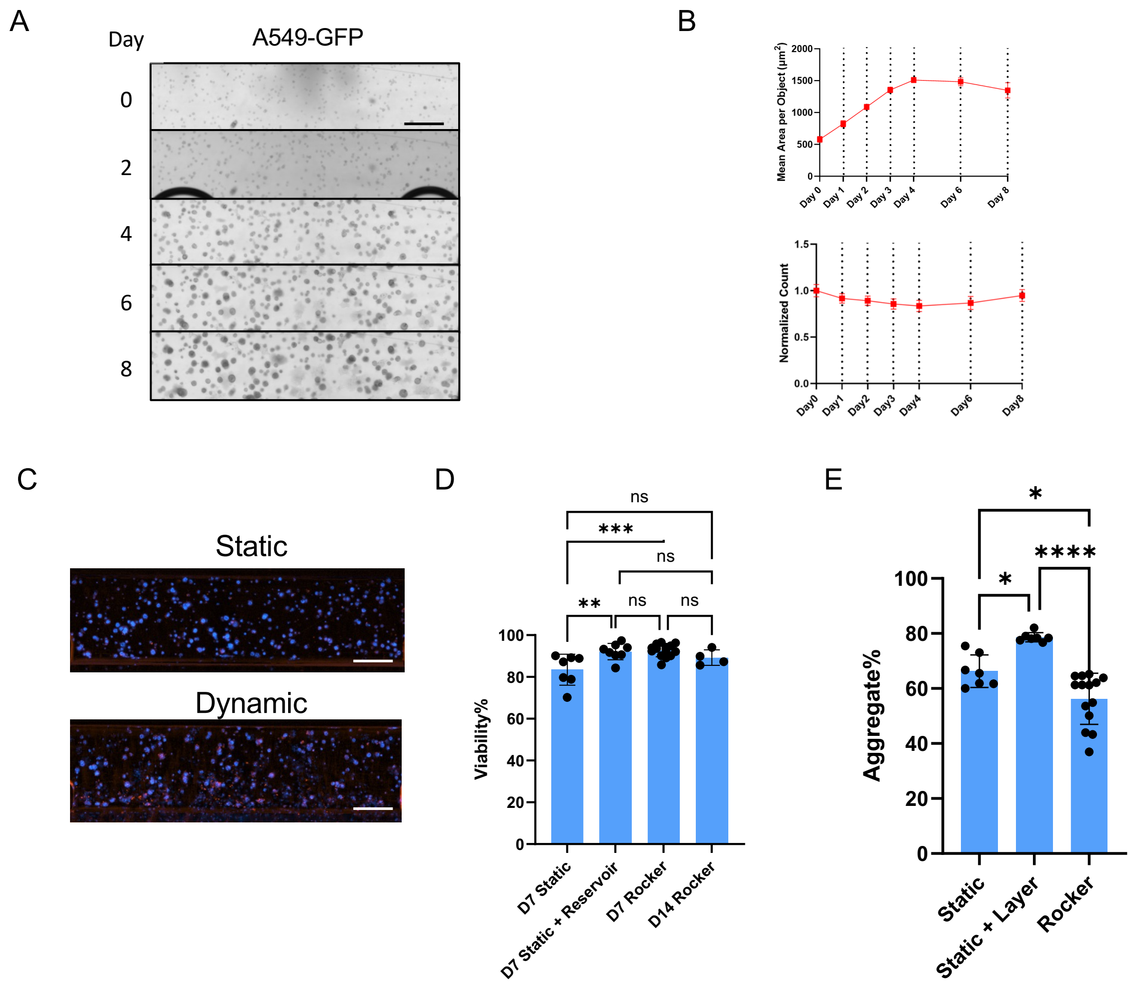

#### Supplementary Figure 6

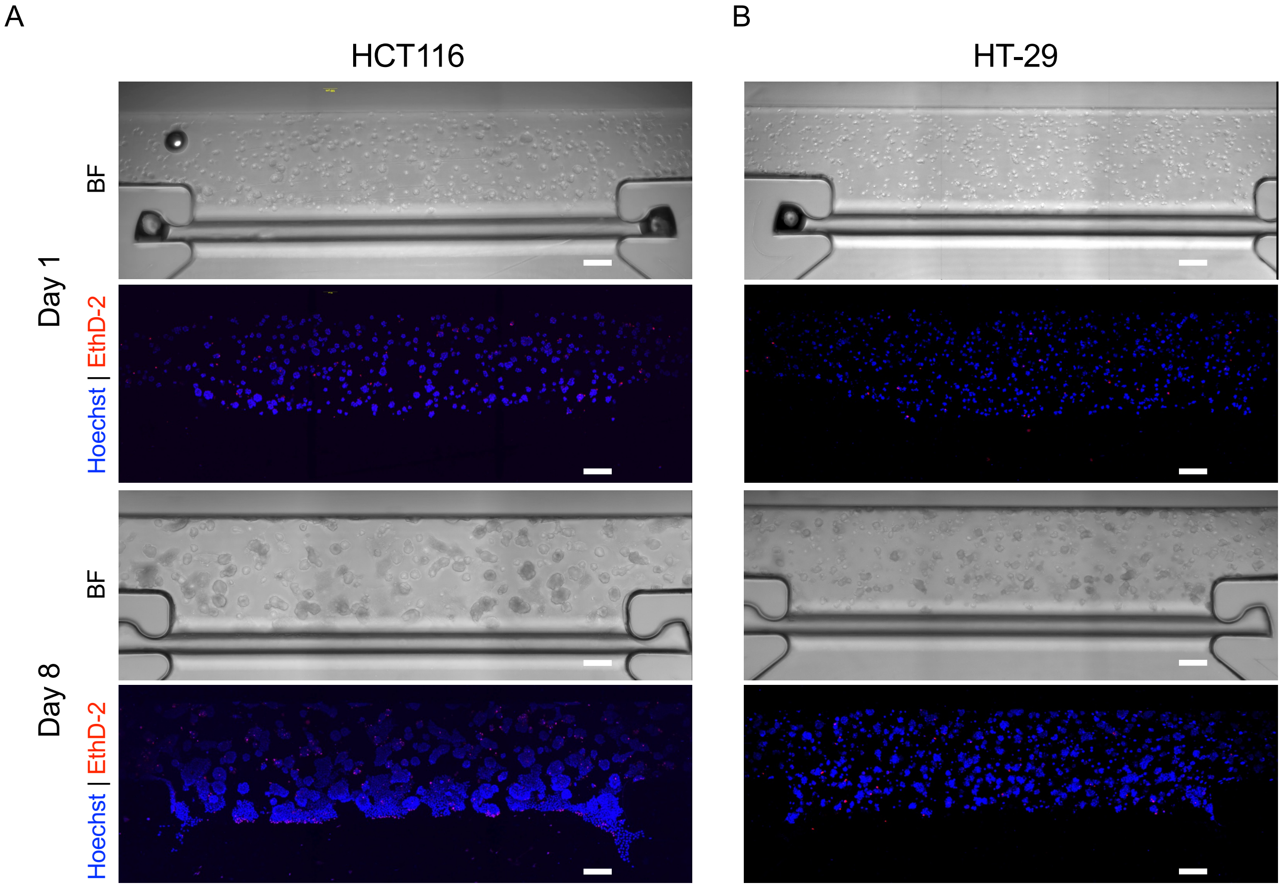

#### Supplementary Figure 7

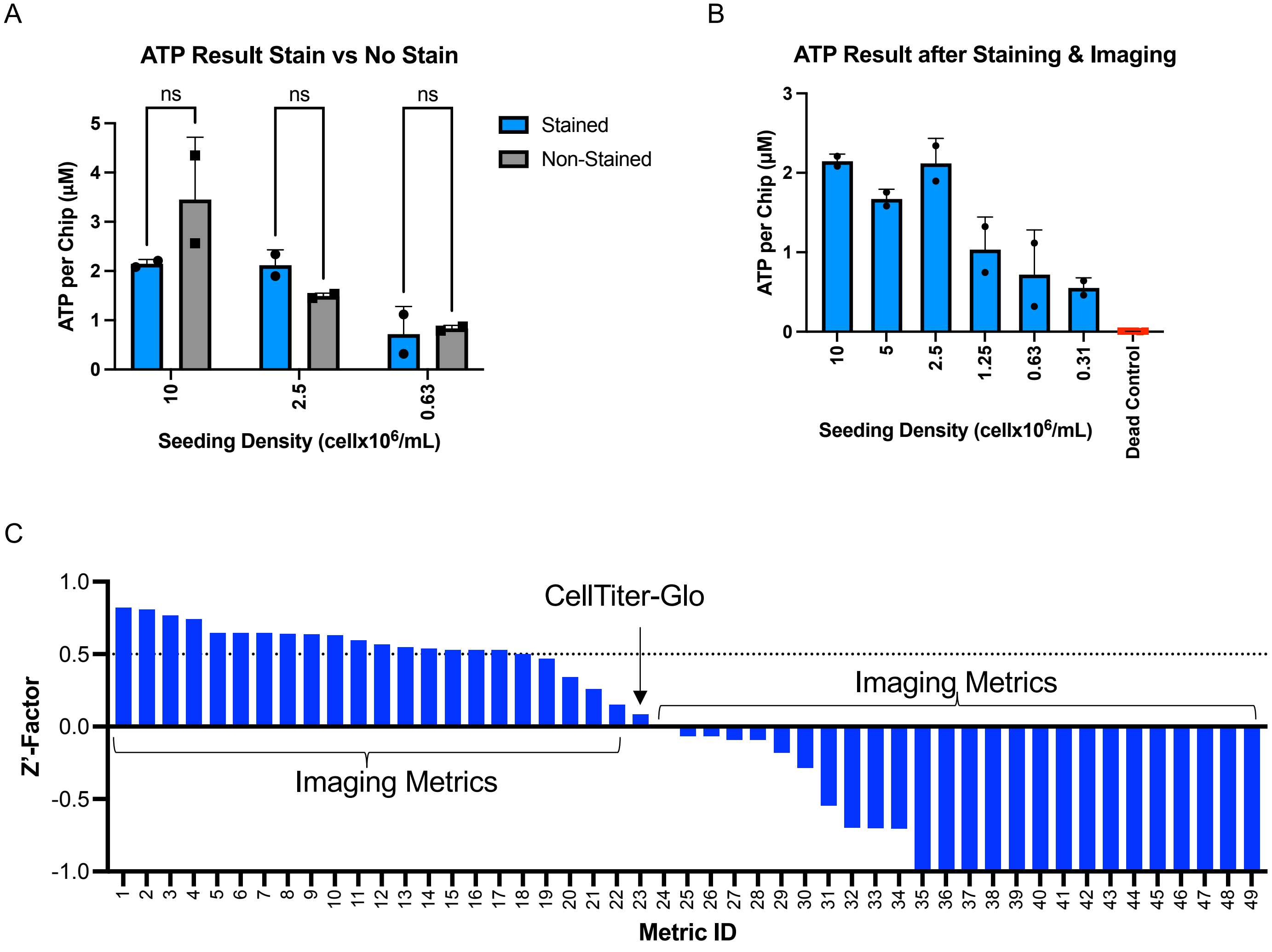

### Supplementary Figure 8

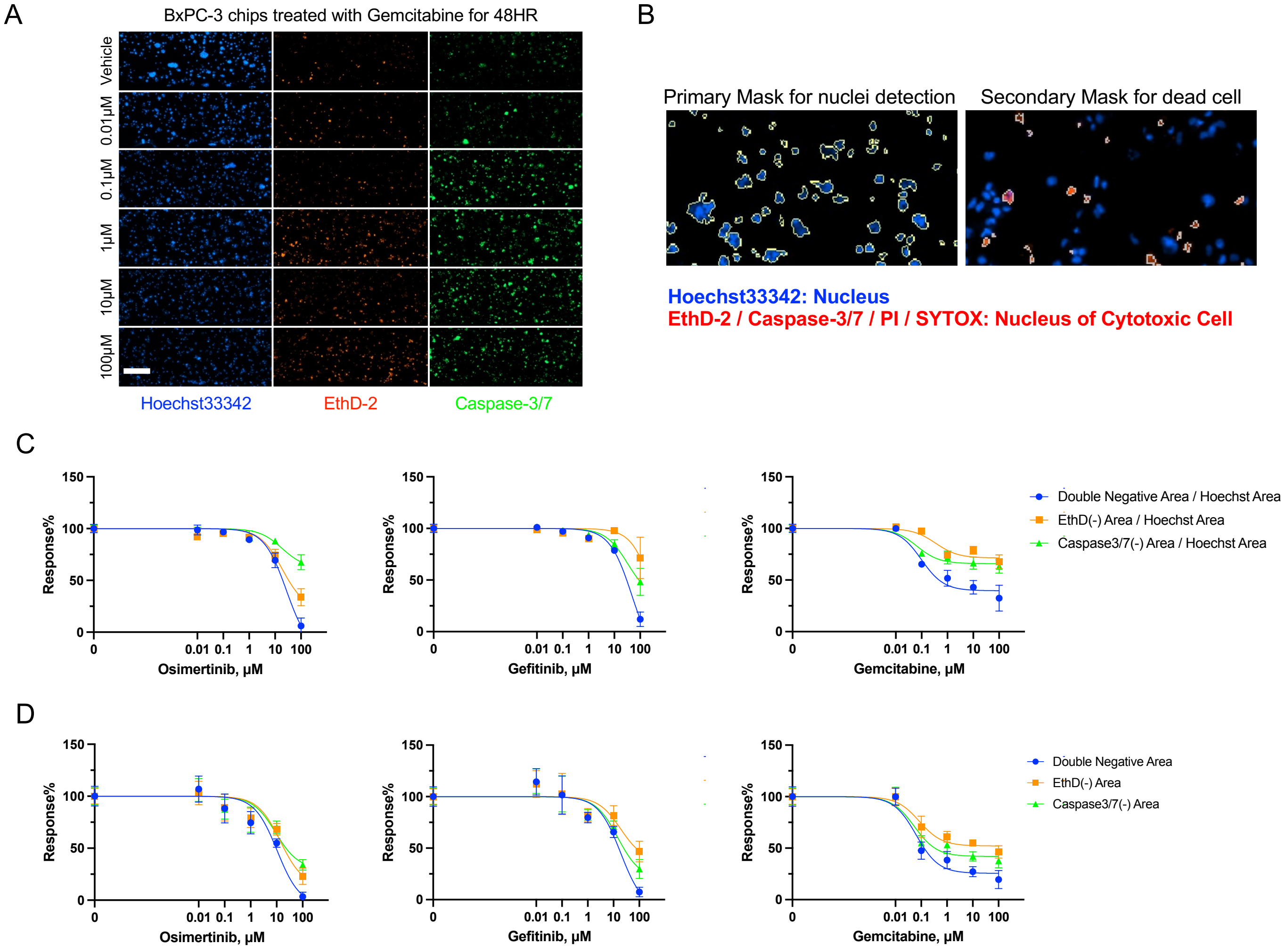

### Supplementary Figure 9

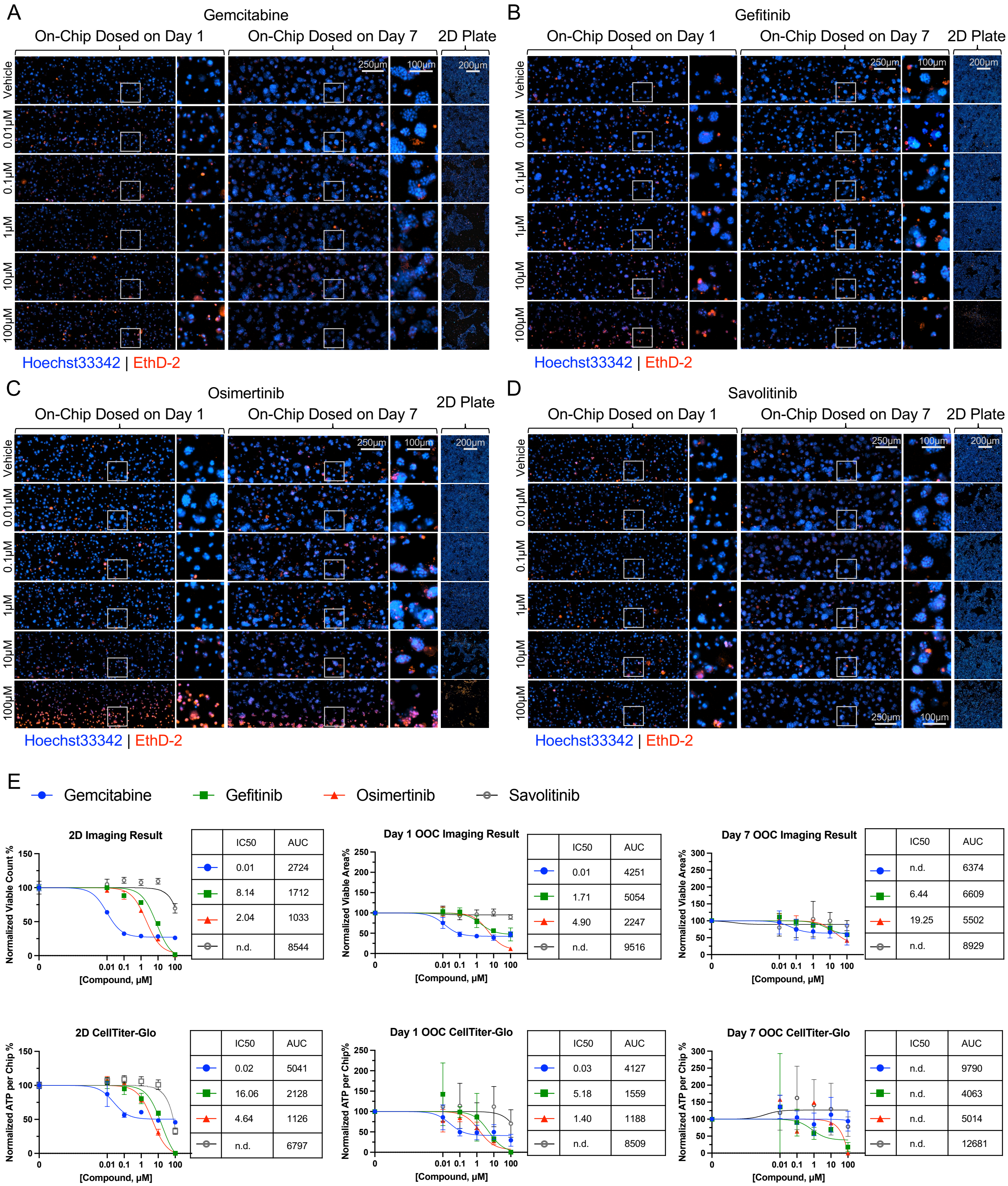


2D Imaging Result

| | IC50 | AUC |
|---|---|---|
| Gemcitabine | 0.01 | 2724 |
| Gefitinib | 8.14 | 1712 |
| Osimertinib | 2.04 | 1033 |
| Savolitinib | n.d. | 8544 |

Day 1 OOC Imaging Result

| | IC50 | AUC |
|---|---|---|
| Gemcitabine | 0.01 | 4251 |
| Gefitinib | 1.71 | 5054 |
| Osimertinib | 4.90 | 2247 |
| Savolitinib | n.d. | 9516 |

Day 7 OOC Imaging Result

| | IC50 | AUC |
|---|---|---|
| Gemcitabine | n.d. | 6374 |
| Gefitinib | 6.44 | 6609 |
| Osimertinib | 19.25 | 5502 |
| Savolitinib | n.d. | 8929 |

2D CellTiter-Glo

| | IC50 | AUC |
|---|---|---|
| Gemcitabine | 0.02 | 5041 |
| Gefitinib | 16.06 | 2128 |
| Osimertinib | 4.64 | 1126 |
| Savolitinib | n.d. | 6797 |

Day 1 OOC CellTiter-Glo

| | IC50 | AUC |
|---|---|---|
| Gemcitabine | 0.03 | 4127 |
| Gefitinib | 5.18 | 1559 |
| Osimertinib | 1.40 | 1188 |
| Savolitinib | n.d. | 8509 |

Day 7 OOC CellTiter-Glo

| | IC50 | AUC |
|---|---|---|
| Gemcitabine | n.d. | 9790 |
| Gefitinib | n.d. | 4063 |
| Osimertinib | n.d. | 5014 |
| Savolitinib | n.d. | 12681 |

### Supplementary Figure 10

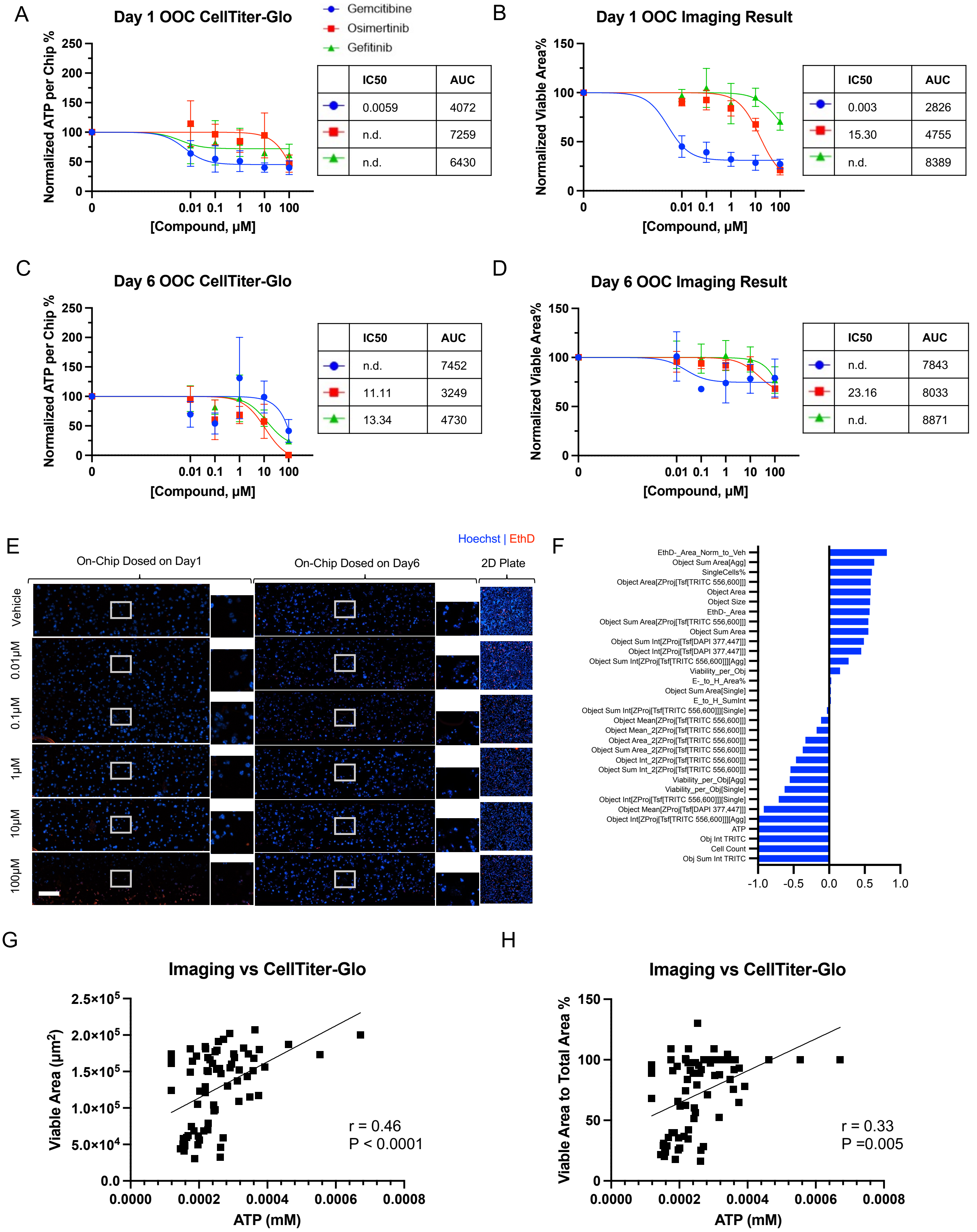

### Supplementary Figure 11

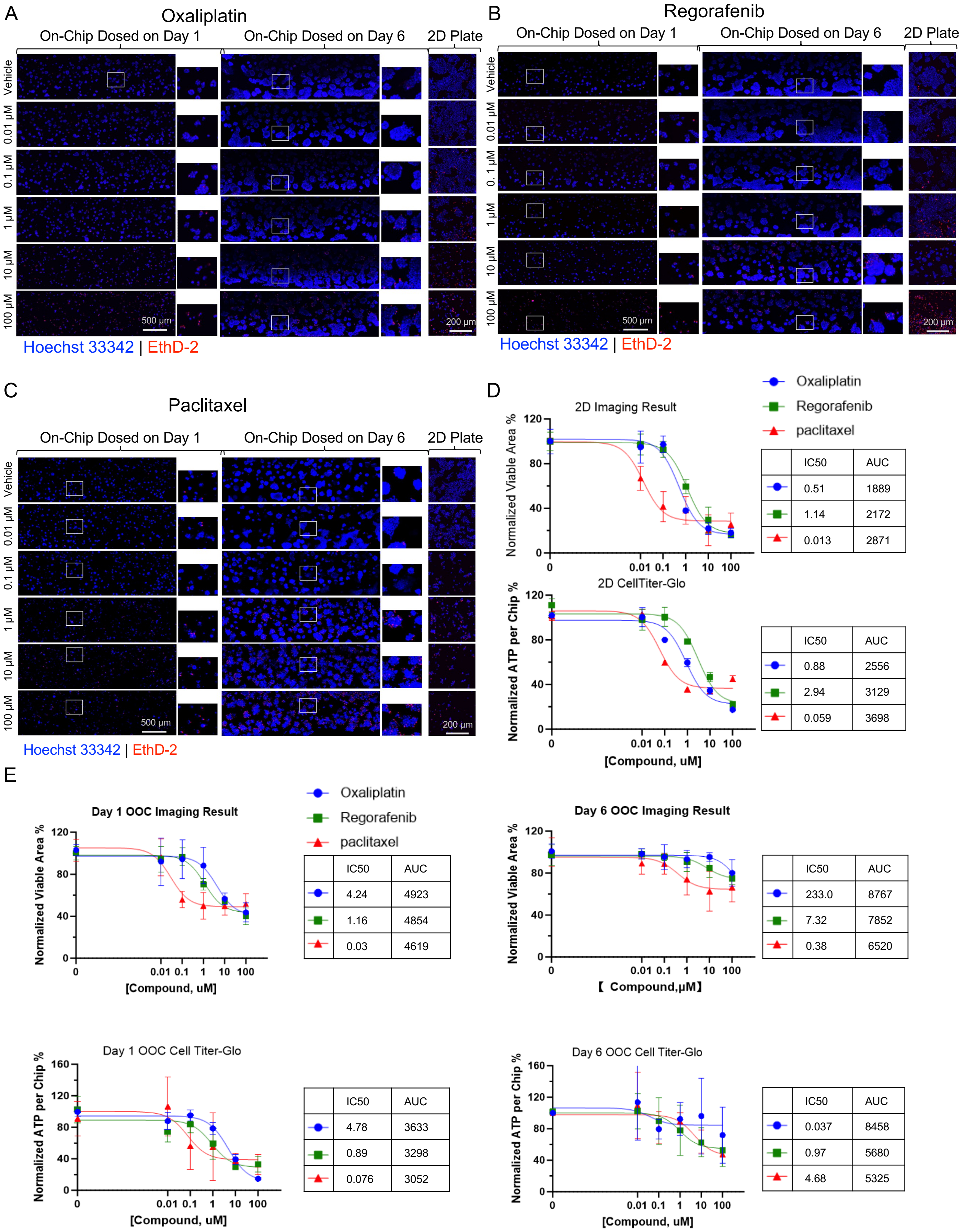

### Supplementary Figure 12

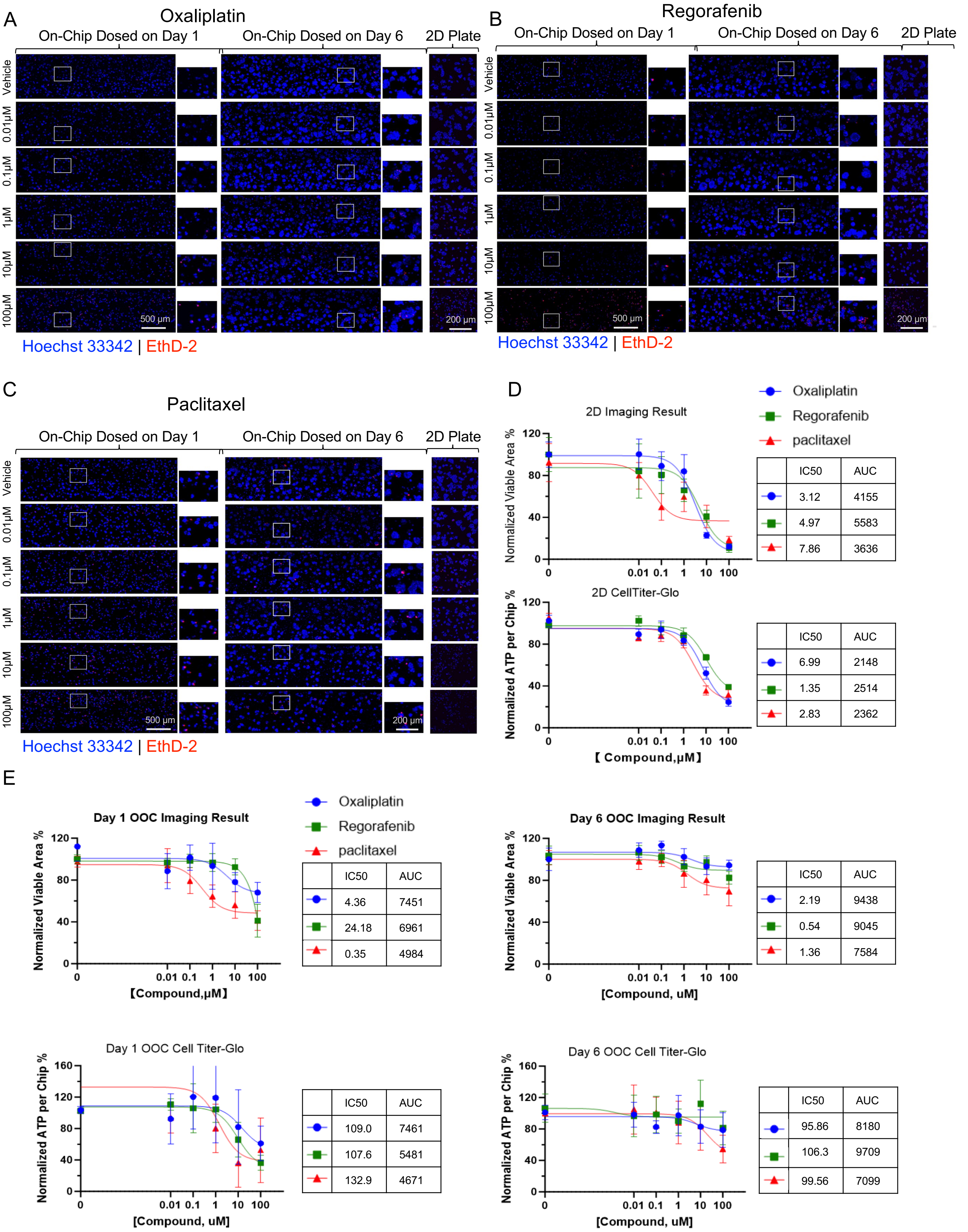


D – 2D Imaging Result

| | IC50 | AUC |
|---|---|---|
| ● | 3.12 | 4155 |
| ■ | 4.97 | 5583 |
| ▲ | 7.86 | 3636 |

D – 2D CellTiter-Glo

| | IC50 | AUC |
|---|---|---|
| ● | 6.99 | 2148 |
| ■ | 1.35 | 2514 |
| ▲ | 2.83 | 2362 |

E – Day 1 OOC Imaging Result

| | IC50 | AUC |
|---|---|---|
| ● | 4.36 | 7451 |
| ■ | 24.18 | 6961 |
| ▲ | 0.35 | 4984 |

E – Day 6 OOC Imaging Result

| | IC50 | AUC |
|---|---|---|
| ● | 2.19 | 9438 |
| ■ | 0.54 | 9045 |
| ▲ | 1.36 | 7584 |

E – Day 1 OOC Cell Titer-Glo

| | IC50 | AUC |
|---|---|---|
| ● | 109.0 | 7461 |
| ■ | 107.6 | 5481 |
| ▲ | 132.9 | 4671 |

E – Day 6 OOC Cell Titer-Glo

| | IC50 | AUC |
|---|---|---|
| ● | 95.86 | 8180 |
| ■ | 106.3 | 9709 |
| ▲ | 99.56 | 7099 |

### Supplementary Figure 13

A

Bright Field overlayed with **original** DAPI Staining mask | Bright Field overlayed with **Predicted** DAPI Staining mask

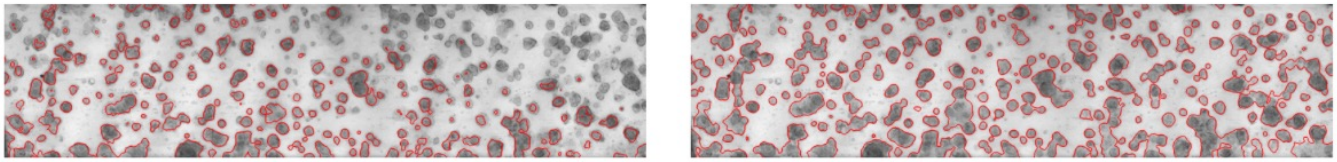

B

DAPI | TRITC | Live

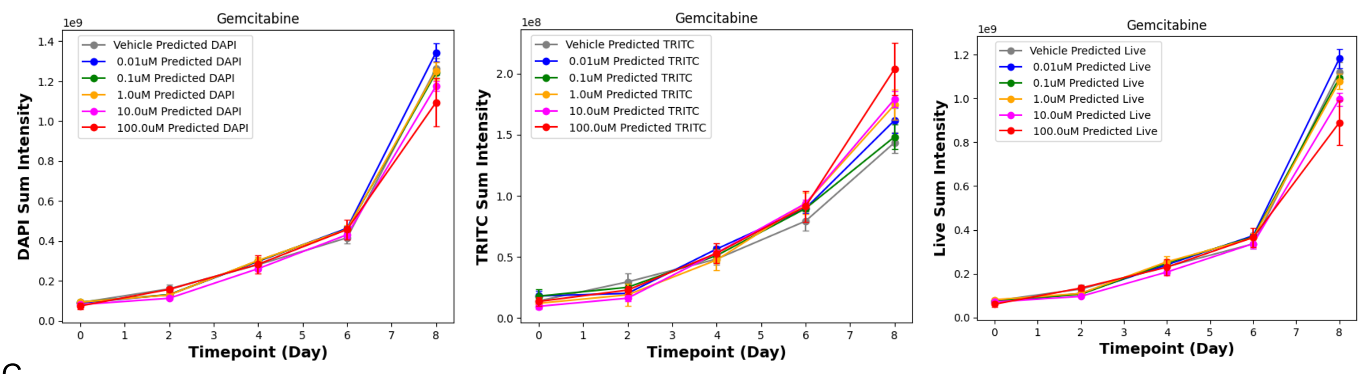


C

DAPI | TRITC | Live

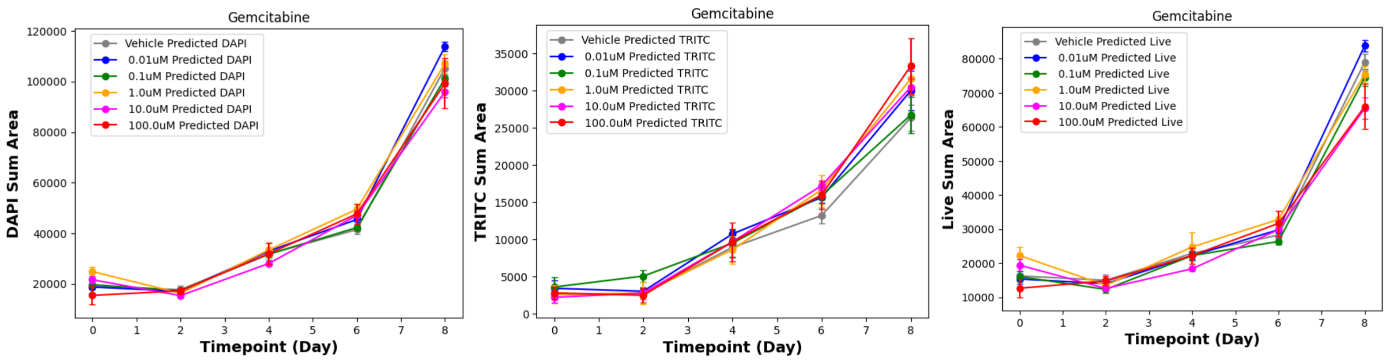
